# Targeted engagement of β-catenin-Ikaros complexes in refractory B-cell malignancies

**DOI:** 10.1101/2023.03.13.532152

**Authors:** Kadriye Nehir Cosgun, Huda Jumaa, Mark E. Robinson, Klaus M. Kistner, Liang Xu, Gang Xiao, Lai N. Chan, Jaewoong Lee, Kohei Kume, Etienne Leveille, David Fonseca-Arce, Dhruv Khanduja, Han Leng Ng, Niklas Feldhahn, Joo Song, Wing-Chung Chan, Jianjun Chen, M. Mark Taketo, Shalin Kothari, Matthew S. Davids, Hilde Schjerven, Julia Jellusova, Markus Müschen

## Abstract

In most cell types, nuclear β-catenin functions as prominent oncogenic driver and pairs with TCF7-family factors for transcriptional *activation* of MYC. Surprisingly, B-lymphoid malignancies not only lacked expression and activating lesions of β-catenin but critically depended on GSK3β for effective β-catenin degradation. Our interactome studies in B-lymphoid tumors revealed that β-catenin formed repressive complexes with lymphoid-specific Ikaros factors at the expense of TCF7. Instead of MYC-activation, β-catenin was essential to enable Ikaros-mediated recruitment of nucleosome remodeling and deacetylation (NuRD) complexes for transcriptional *repression* of MYC.

To leverage this previously unrecognized vulnerability of B-cell-specific repressive β-catenin-Ikaros-complexes in refractory B-cell malignancies, we examined GSK3β small molecule inhibitors to subvert β-catenin degradation. Clinically approved GSK3β-inhibitors that achieved favorable safety prof les at micromolar concentrations in clinical trials for neurological disorders and solid tumors were effective at low nanomolar concentrations in B-cell malignancies, induced massive accumulation of β-catenin, repression of MYC and acute cell death. Preclinical *in vivo* treatment experiments in patient-derived xenografts validated small molecule GSK3β-inhibitors for targeted engagement of lymphoid-specific β-catenin-Ikaros complexes as a novel strategy to overcome conventional mechanisms of drug-resistance in refractory malignancies.

**HIGHLIGHTS:**

- Unlike other cell lineages, B-cells express nuclear β-catenin protein at low baseline levels and depend on GSK3β for its degradation.
- In B-cells, β-catenin forms unique complexes with lymphoid-specific Ikaros factors and is required for Ikaros-mediated tumor suppression and assembly of repressive NuRD complexes.
- CRISPR-based knockin mutation of a single Ikaros-binding motif in a lymphoid *MYC* superenhancer region reversed β-catenin-dependent Myc repression and induction of cell death.
- The discovery of GSK3β-dependent degradation of β-catenin as unique B-lymphoid vulnerability provides a rationale to repurpose clinically approved GSK3β-inhibitors for the treatment of refractory B-cell malignancies.

**GRAPHICAL ABSTRACT:** 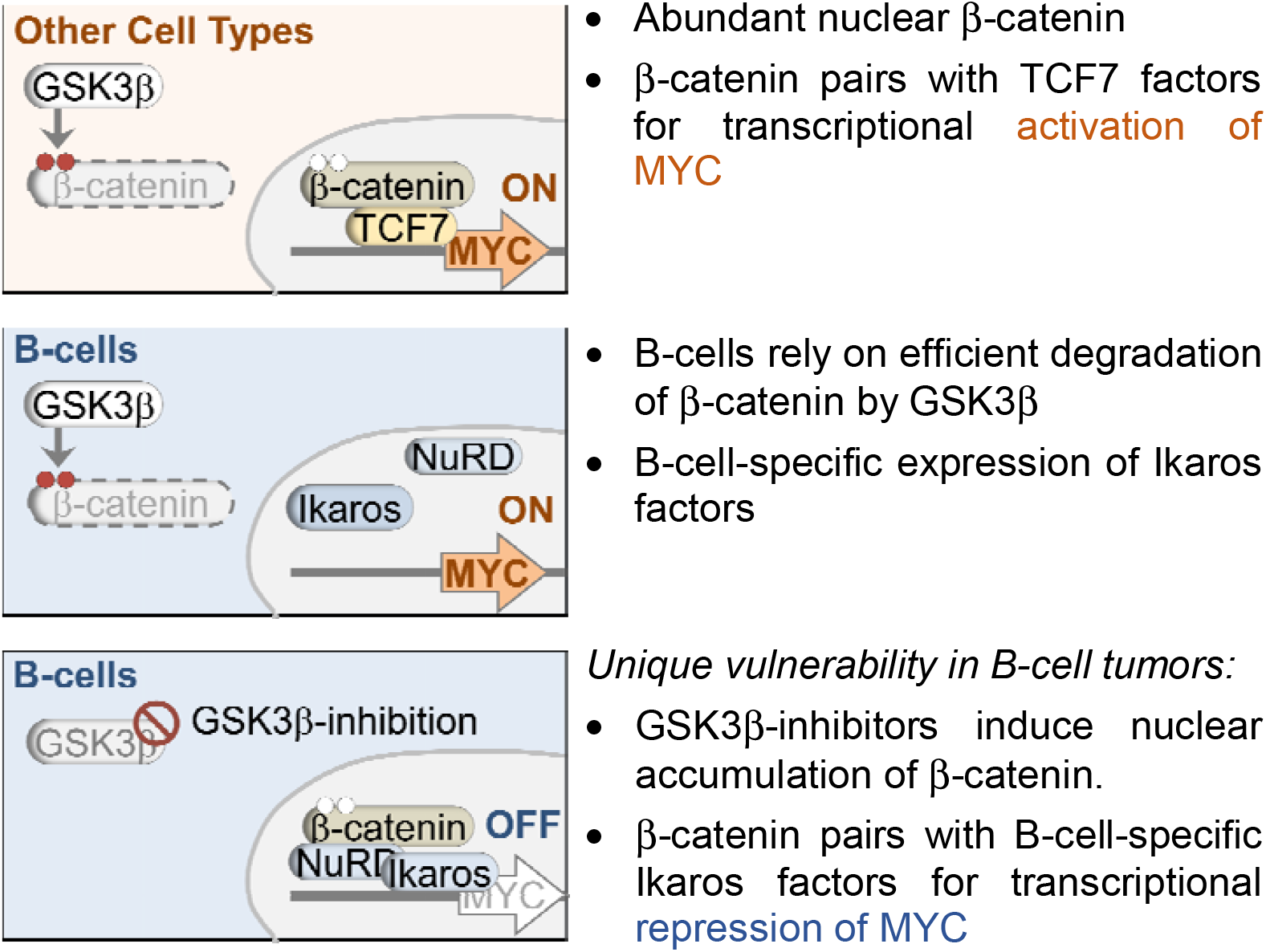

- Abundant nuclear β-catenin
- β-catenin pairs with TCF7 factors for transcriptional activation of MYC
- B-cells rely on efficient degradation of β-catenin by GSK3β
- B-cell-specific expression of Ikaros factors

*Unique vulnerability in B-cell tumors:*

- GSK3β-inhibitors induce nuclear accumulation of β-catenin.
- β-catenin pairs with B-cell-specific Ikaros factors for transcriptional repression of MYC

## INTRODUCTION

The WNT/β-catenin pathway is involved in fundamental processes including embryonic development, organogenesis, and tissue homeostasis^1-6^. β-catenin protein levels are tightly regulated by β-catenin-degradation, which is initiated by Glycogen Synthase Kinase 3β (GSK3β) and the scaffolding proteins Axin1, Axin2 and Adenomatous Polyposis Coli (APC)^7-9^. In the absence of Wnt ligands, β-catenin is phosphorylated by GSK3β on N-terminal serine and threonine residues encoded by exon 3 for subsequent proteasomal degradation^8-9^. Conversely, Wnt ligands stabilize β-catenin and induce its nuclear accumulation to promote transcription of Wnt target genes including MYC^4-6,9^. β-catenin functions as a central driver of MYC-expression, proliferation, and survival in multiple epithelial, neuronal, and mesenchymal lineages^4-6^, but is dispensable for hematopoietic development^10-12^. Intermediate levels of β-catenin signaling were shown to promote survival and proliferation of hematopoietic stem and progenitor cells^13^, as well as multiple stages of T-cell development^14-16^. Targeted removal of GSK3β-phosphorylation sites and β-catenin accumulation in *Ctnnb1*^ex3fl/+^ mice, had detrimental effects on hematopoiesis^17-18^, and suppressed early T-cell development^19^. This contrasts with studies in colon cancer, melanoma, and other epithelial cancers, where genetic accumulation of β-catenin results in acceleration of proliferation and malignant transformation^2,20-21^. The role of β-catenin signaling in myeloid and T-cell malignancies is controversial: While earlier studies demonstrated an important role of β-catenin in the initiation of myeloid (AML, CML)^22-24^ and T-cell leukemia (T-ALL)^25-26^, a new genetic mouse model provided evidence that T-cell development and AML leukemia-initiation do not require β-catenin^12^. β-catenin lacks a DNA binding domain and interacts with TCF7-family transcription factors^2-3^ in epithelial, mesenchymal, neuronal, and myeloid cells to induce transcriptional activation of WNT-target genes including MYC^4-6^. Here we show that genetic and pharmacological β-catenin accumulation selectively impact B-lymphoid cells by transcriptional repression of MYC. This unexpected outcome of β-catenin signaling in B-lymphocytes was predicated on previously unrecognized repressive complexes between β-catenin and lymphoid-specific Ikaros (IKZF1 and IKZF3) transcription factors.

## RESULTS

### Lack of β-catenin signaling in B-cell development and B-lymphoid malignancies

In a computational pan-cancer analysis of oncogenic drivers in eight defined signaling pathways, activating lesions of the β-catenin signaling pathway were strongly selected in 12 cancer types but showed evidence of negative selective pressure in B-cell acute lymphoblastic leukemia (B-ALL) and B-cell lymphoma (**Figure 1a**). Studying individual β-catenin pathway lesions (FATHMM score >0.5) in 66,949 cancer samples encompassing 16 types of solid tumors and hematological malignancies^27^, we found frequent mutations, including of *CTNNB1* itself (1.6%) and its negative regulators *APC* (8.9%), *AXIN1* (1.4%), *AXIN2* (1.1%) and *GSK3B* (0.6%). In contrast, among 2,137 B-cell malignancies, cases with activating β-catenin pathway lesions were markedly underrepresented (expected 264, observed 17, χ² test *P*=1.2 E-12; **Figure 1b, Table S1)**. Since mutation frequencies for two of the five oncogenic drivers of β-catenin signaling were also reduced in T-cell malignancies (*AXIN2, GSK3B*; **Table 1**), we examined whether these differences reflect a general reduction of β-catenin signaling in lymphoid lineages, encompassing B-, T- and NK-cells. To this end, we measured β-catenin signaling in Axin2-turquoise reporter transgenic mice^28^. While β-catenin signaling was clearly detectable in T-cells and NK-cells, this was not the case for B-cells (**Figure 1c**), suggesting that attenuation of β-catenin signaling represents a unique feature of B-lymphocytes. Immunohistochemistry confirmed low baseline levels of β-catenin protein expression in B-lymphoid tissues (n=24) compared to consistently high β-catenin expression in myeloid bone marrow cells, as well as epithelial and mesenchymal cell types (n=30; **Figure 1f, S1a**). Detailed flow cytometry analysis of Axin2-turquoise transgenic mice corroborated that, unlike thymic and mature T-cell development, B-cells consistently lacked β-catenin signaling throughout early and late stages of development (**Figure S1b-c**). Interestingly, β-catenin mRNA levels were comparable between B-cell malignancies and solid tumors (RNA-seq; **Figure 1d**). While β-catenin mRNA levels were similar in B-cells compared to other cell types, β-catenin protein levels (reverse phase protein arrays, proteomics) were markedly suppressed in B-lymphoid cell lines (**Figure 1d**), suggesting that B-cells may be more adept than other cell types at clearing β-catenin protein by GSK3β-mediated degradation^8,9^. Mirroring low β-catenin protein levels, B-cell leukemia and lymphoma cells were resistant to CRISPR-mediated deletion of *CTNNB1*^29^, while solid tumors, in particular gastrointestinal tumors, were sensitive to loss of β-catenin (**Figure 1e**). We confirmed low baseline β-catenin protein expression in a panel of 84 B-cell malignancies compared to consistently high β-catenin protein levels in epithelial cancers (lung, colon, melanoma; n=27) by immunohistochemistry (**Figure S2**). B-cell-specific lack of nuclear β-catenin was further corroborated by nuclear and cytoplasmic cell fractionation and Western blot analysis: Nuclear accumulation of β-catenin was consistently detected at high levels in epithelial tumor cell lines (n=16), including lung and colon cancer and malignant melanoma, but not in B-ALL (n=7) and mature B-cell malignancies (n=15; **Figures 1g, S2**).

**Figure 1:**
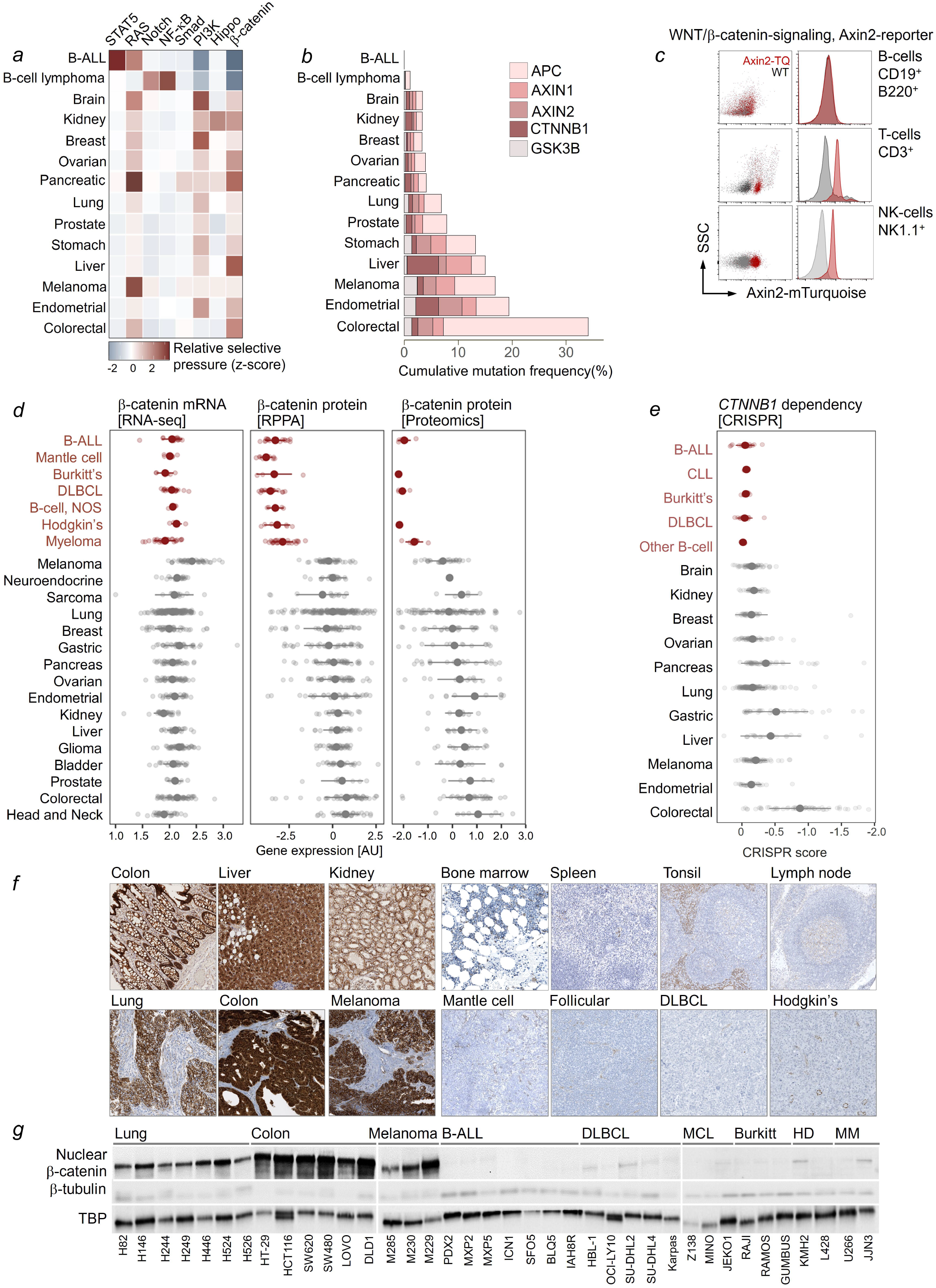
B-lymphoid cells are exempt from β-catenin signaling. (**a**) Computational analyses of positive (red) or negative (blue) selection of known driver mutations along eight signaling pathways were performed for 14 cancer types, including B-ALL and B-cell lymphoma. (**b**) Frequencies of pathogenic mutations (FATHMM score > 0.5) of β-catenin (*CTNNB1*; filtered for hot spot mutations in exon 3), *APC, AXIN1, AXIN2* and GSK3β (*GSK3B*) are depicted for 14 types of cancer including B-cell malignancies and solid tumors. (**c**) Analysis of Wnt/β-catenin activity in B-cells (CD19^+^ B220^+^), T-cells (CD3^+^) and NK-cells (NK.K1^+^) in Axin2-mTurquoise reporter transgenic mice. (**d**) Transcriptional analysis of 1,389 cancer cell lines by RNA-seq for the expression of CTNNB1 (left). Protein levels of β-catenin were assessed by RPPA (middle) and mass-spectrometry (right) in B-cell malignancies compared to solid tumors. (**e**) CTNNB1 dependency among human cancer cell lines evaluated by CRISPR loss-of function screen^29^. (**f**) Representative immunohistochemical stainings for β-catenin (brown) in normal epithelial and lymphoid tissues in comparison to lung cancer (n=15), colon cancer (n=25), malignant melanoma (n=5), mantle cell lymphoma (MCL; n=26), follicular lymphoma (n=38), diffuse large B-cell lymphoma (DLBCL; n=35) and Hodgkin’s lymphoma (HD; n=44) using H&E as counterstain. (**g**) Western blot analysis for β-catenin, β-tubulin, and TATA box binding protein (TBP) on nuclear fractions of lung and colon cancer, malignant melanoma, B cell acute lymphoblastic leukemia (B-ALL), DLBCL, MCL, Burkitt’s lymphoma, HD and multiple myeloma cell lines. Western blots of the cytoplasmic fractions from the same cell lysates are shown in **Figure S2**.

### B-lymphoid cells critically depend on GSK3β-mediated degradation of β-catenin protein

To follow up on the observation that B-cells consistently lacked expression and activity of β-catenin protein despite relatively high mRNA levels, we studied GSK3β-dependent protein degradation of β-catenin in B-lymphoid cells. To this end, we crossed *Ctnnb1*^ex3fl/+^ mice^30^ with a B-cell-specific *Mb1*-Cre deleter strain. Expression of Cre in this model recapitulates β-catenin accumulation as the common outcome of cancer-associated genetic lesions in this pathway, that are commonly found throughout the entire spectrum of cancer but not in B-lymphoid malignancies (**Figure 1b**; **Table S1**). Cre-mediated removal of GSK3β-phosphorylation sites of β-catenin abrogated GSK3β-mediated degradation, resulting in β-catenin accumulation from earliest stages of B-cell development. While pro-B and pre-BI cells (Hardy fractions A-C) tolerated β-catenin accumulation, B-lymphopoiesis beyond pre-BCR^+^ stages (Hardy fractions C’-F and mature B-cells) of development was profoundly suppressed *in vivo* **(Figures 2a-b, S3**). To model defective GSK3β-mediated degradation of β-catenin in common subtypes of B-ALL, we transduced B-cell precursors from *Ctnnb1*^ex3fl/+^ mice with BCR-ABL1 and NRAS^G12D^ oncogenes and tamoxifen-inducible Cre (Cre-ER^T2^) or empty vector control (ER^T2^). Inducible β-catenin accumulation subverted competitive fitness of B-ALL cells, abolished colony formation and induced G_0_/G_1_ phase cell cycle arrest (**Figure 2c-e**). Transplant experiments revealed that β-catenin stabilization in BCR-ABL1-driven B-ALL cells subverted leukemia-initiation *in vivo*: Compared to controls, β-catenin stabilization reduced the frequency of leukemia-initiating cells by about 40-fold (**Figure 2f-g**).

**Figure 2:**
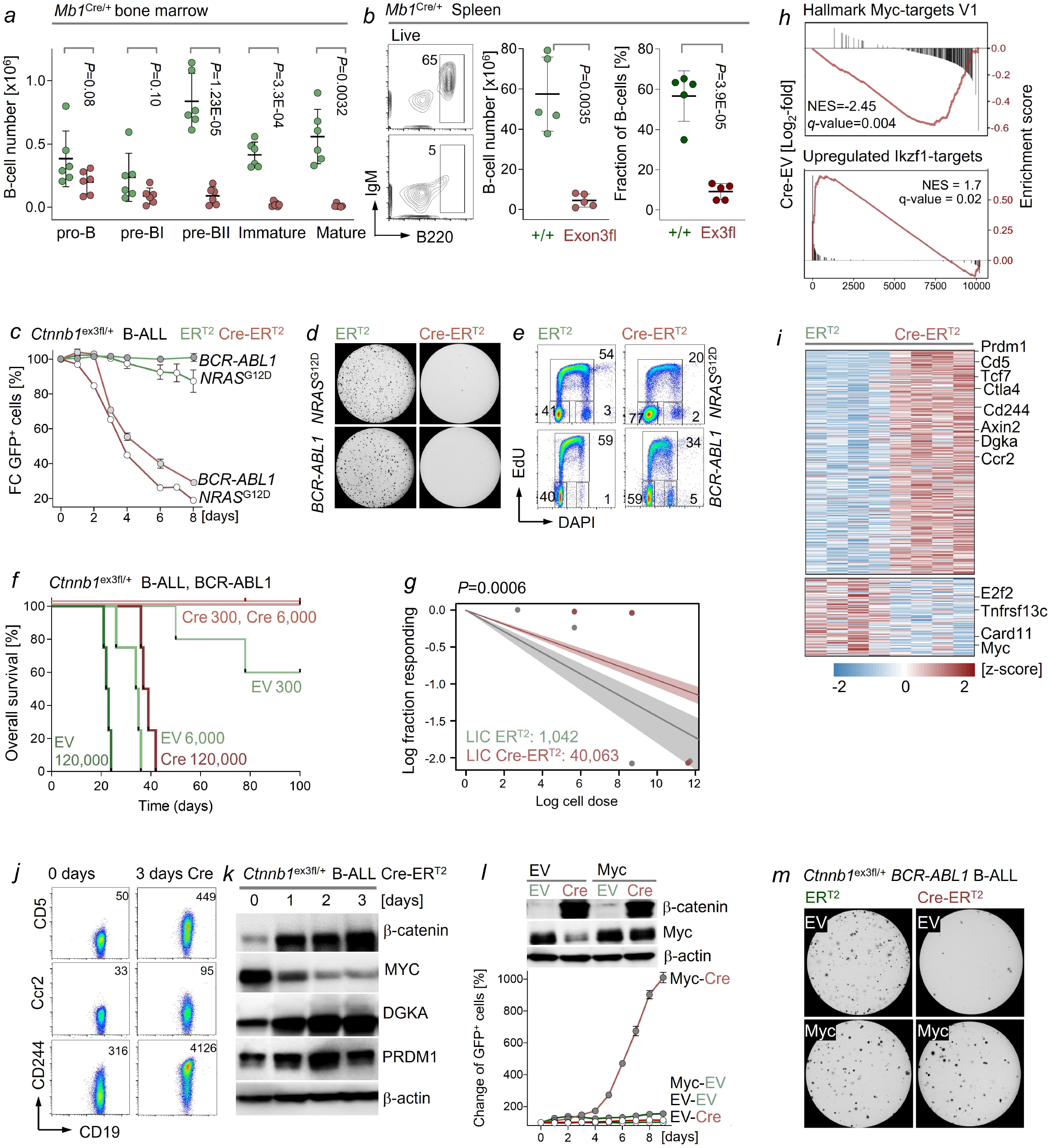
Genetic accumulation of β-catenin suppresses B-cell development and malignant transformation in vivo. (**a-b**) B-cell development in the bone marrow and spleen of *Mb1*^Cre/+^ *Ctnnb1*^ex3fl^ mice was analyzed by flow cytometry. (**a**) The numbers of pro-B cells (CD43^+^ B220^low^ IgM^-^ BP1^-^) and pre-BI cells (CD43^+^ B220^low^ IgM^-^ BP1^+^), pre-BII cells (CD43^-^ B220^low^ IgM^-^), immature B cells (CD43^-^ B220^low^ IgM^+^) and mature B cells (CD43^-^ B220^high^ IgM^+^) in the bone marrow of *Mb1*^Cre/+^ *Ctnnb1*^ex3fl^ mice are shown from 6 independent experiments. (**b**) Absolute numbers and frequencies of B220^+^ splenic B-cells and representative FACS plots are shown. *Ctnnb1*^ex3fl/+^ *BCR*-*ABL1 or NRAS*^G12D^ transformed B-ALL cells were transduced with vectors expressing GFP and 4-hydroxy-tamoxifen (4-OHT)-inducible Cre (Cre-ER^T2^) or ER^T2^. (**c**) Changes of percentages of GFP^+^ cells were monitored for 8 days following 4-OHT addition, data representative of three independent experiments (triplicates). (**d**) B-ALL cells were sorted for GFP expression and plated for colony formation assays after 4-OHT treatment. Representative images for 10 days after plating. (**e**) Cell cycle phases of *Ctnnb1*^ex3fl/+^ *NRAS*^G12D^ or *BCR-ABL1* B-ALL cells were measured by EdU incorporation in combination with DAPI staining 2 days after β-catenin accumulation. Data shown are representative of two independent experiments (triplicates). (**f-g**) Extreme limiting dilution analysis (ELDA) was performed to assess effects of β-catenin accumulation on leukemia-initiation capacity (LIC) of *BCR-ABL1*-driven B-ALL cells. (**f**) Kaplan-Meier analysis of overall survival in each group and dose level (n=4; *P*=8.2E-10; Log-rank test). (**g**) LIC was determined in B-ALL cells with β-catenin accumulation (1 in 40,063 cells) and control cells (1 in 1,042; *P*=0.0006). (**h-i**) Gene expression changes were studied by RNA-seq analysis in *Ctnnb1*^ex3fl/+^ B-ALL one day after 4-OHT-treatment (n=4). (**h**) Gene set enrichment analysis (GSEA) identified depletion of Myc target genes and enrichment of Ikaros target genes as top-ranking gene sets following β-catenin accumulation. (**i**) Genes that were upregulated (n=354) or down-regulated (n=119) upon β-catenin stabilization are shown as heatmap. (**j**) Flow cytometry analysis to validate CD5, Ccr2 and CD244 (2B4) upregulation 3 days after Cre-mediated stabilization of β-catenin. (**k**) Changes in protein levels of β-catenin, Myc, Dgka, Prdm1 were studied by Western blot 0-3 days after β-catenin activation. *BCR-ABL1* transformed *Ctnnb1*^ex3fl/+^ B-ALL cells expressing Cre-ER^T2^ or ER^T2^ (puromycin selected) were transduced with GFP-tagged Myc or empty vector (EV). **(l)** Expression of β-catenin and Myc in FACS-sorted GFP^+^ cells was confirmed by Western blot 3 days after 4-OHT treatment. FACS analyses were performed to monitor enrichment or depletion of GFP^+^ cells (Myc vs EV) upon β-catenin activation. Representative data from three independent experiments (triplicates) is shown. (**m**) Colony formation ability of cells expressing Myc, or empty vector (EV) was assessed 2 days after 4-OHT induced β-catenin accumulation. Data shown is a representative of two independent experiments (triplicates).

### β-catenin-accumulation in B-lymphoid cells results in transcriptional repression of MYC

RNA-seq analysis of β-catenin-dependent gene expression changes revealed enrichment for two principal gene sets, namely suppression of Myc-targets and of activation of transcriptional targets of the Ikaros zinc finger protein IKZF1 (**Figure 2h**). While Myc and E2f were repressed, molecules related to B-cell anergy (Prdm1, Cd5, Dgka, Cd244, Ctla4), and β-catenin signaling (Tcf7, Axin2) were strongly upregulated upon β-catenin-activation (**Figure 2i-k**). β-catenin-mediated repression of Myc in murine B-ALL cells was in striking contrast to previous findings of *MYC* as a classical target of β-catenin-mediated transcriptional activation in epithelial cells^4-6^. Consistent with gene set enrichment analyses (**Figure 2h**), genetic rescue experiments identified suppression of Myc as central mechanistic element of β-catenin-mediated cell death in mouse B-ALL cells: Reconstitution of MYC expression rescued the deleterious effects of β-catenin-accumulation and restored both colony formation and competitive fitness of B-ALL cells (**Figure 2l-m**).

To determine functional consequences of β-catenin accumulation in human cells, we studied doxycycline-inducible expression of stabilized β-catenin in B-lymphoid (B-ALL, mantle cell, Burkitt’s lymphoma, DLBCL; n=13) cell lines and patient-derived xenografts (PDX), myeloid leukemia (n=4), and colon and lung epithelial cell lines (n=6). In B-lymphoid cells, inducible β-catenin accumulation suppressed MYC-expression, compromised clonal fitness, colony formation, cell proliferation and induced cell death (**Figure 3a-f**). Inducible β-catenin accumulation had no significant effects in colon and lung cells and increased competitive fitness and colony formation in myeloid leukemia cells (**Figure 3a-e**). Together, these results suggest that B-lymphoid cells fundamentally differ from myeloid and epithelial cell types in that they are not permissive to accumulation of β-catenin.

**Figure 3:**
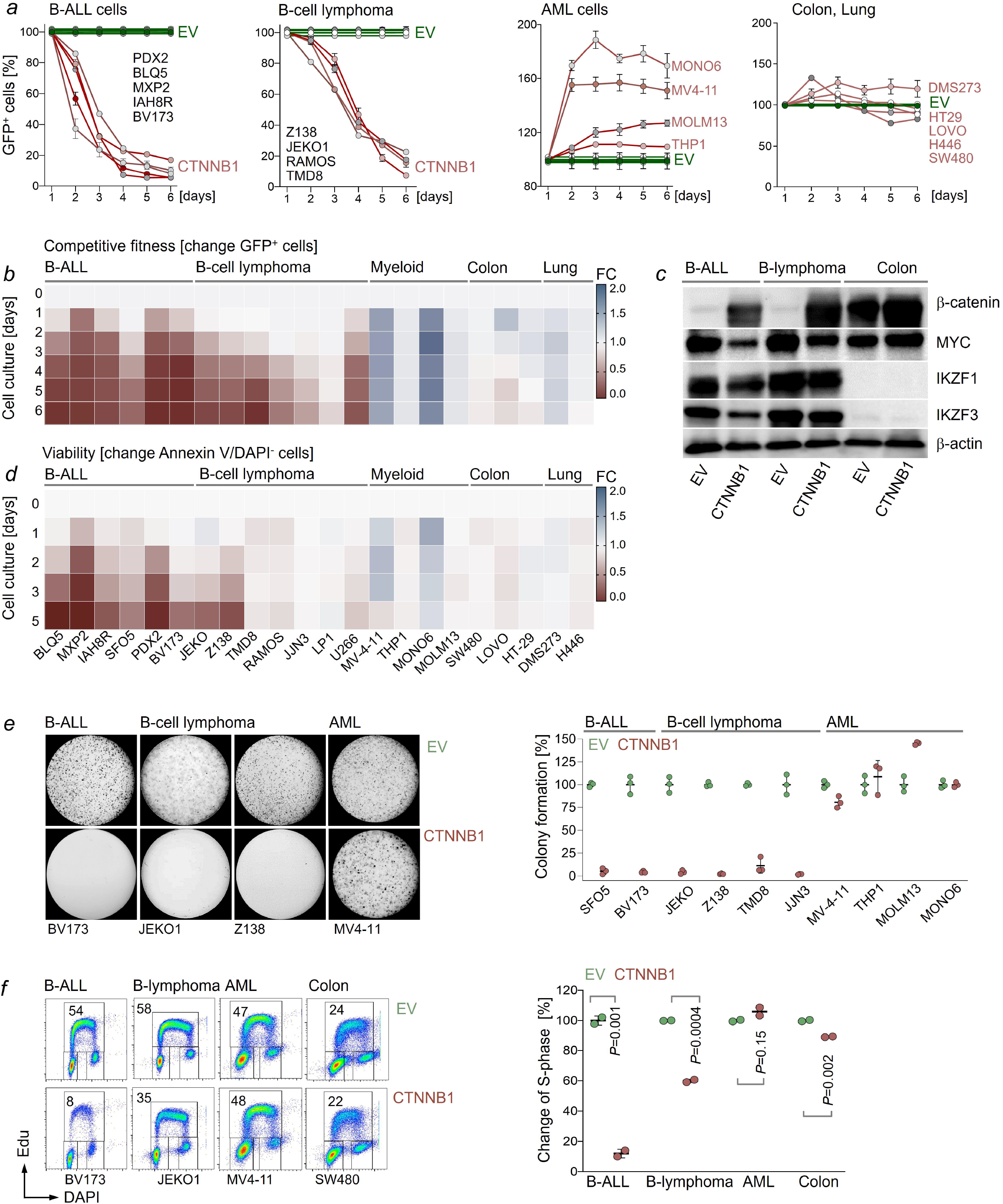
Deleterious effects of β-catenin-accumulation in B-lymphoid but not myeloid and epithelial cells. Human cancer cell lines or patient derived xenografts were transduced with Tet-3G doxycycline-inducible vectors for expression of GFP tagged stabilized β-catenin with point mutations of GSK3β-phosphorylation sites (CTNNB1) or empty vector (EV). (**a-b**) Doxycycline was added to induce expression of β-catenin and GFP and changes in the percentages of GFP^+^ cells were monitored by FACS. Data shown are representative of two independent experiments (triplicates). (**c**) B-ALL (BLQ5), mantle cell lymphoma (JEKO1), and colon cancer (LOVO) cell lines carrying inducible β-catenin constructs were treated with doxycycline for two days. Western blot was performed to detect the expression of β-catenin, MYC and the lymphoid transcription factors IKZF1 and IKZF3, using β-actin as loading control. (**d**) Cell viability following β-catenin accumulation was monitored over time by flow cytometry based on Annexin V and DAPI staining. Data shown are representative of two independent experiments. (**e**) One day after doxycycline treatment GFP^+^ cells were FACS sorted (99.8% pure) and plated on methylcellulose medium for colony forming assays. Colonies were imaged and counted 14 days after plating. Representative images from two independent experiments are shown (triplicates). Cell lines left to right: BV173 (B-ALL), JEKO and Z138 (mantle cell lymphoma), and MV4-11 (acute myeloid leukemia). (**f**) Cell cycle analyses were performed by measuring EdU incorporation 2 days after doxycycline mediated expression of stabilized β-catenin. Changes in frequencies of cells in S phase following β-catenin accumulation were shown. Data are representative of two independent experiments (two replicates each). Cell lines left to right: BV173 (B-ALL), JEKO (mantle cell lymphoma), MV4-11 (acute myeloid leukemia) and SW480 (colon cancer).

### β-catenin forms repressive complexes with Ikaros factors and NuRD components in B-lymphoid cells

β-catenin lacks a DNA binding domain and interacts with TCF7-family transcription factors^2-3^ in epithelial, mesenchymal, neuronal, and myeloid cells to induce transcriptional activation of MYC^4-6^. Given that β-catenin-accumulation unexpectedly repressed MYC in B-lymphoid cells, we systematically compared β-catenin-interacting proteins in B-lymphoid, myeloid, and epithelial cell types. In an initial experiment, we preformed co-immunoprecipitation (Co-IP) experiments and identified β-catenin binding proteins in murine B-ALL cells by mass-spectrometry. Besides known β-catenin interacting proteins (Apc, Axin1, Gsk3β)^7-8^, the proteins with the highest enrichment of binding to β-catenin included the lymphoid-specific Ikaros transcription factors Ikaros (*Ikzf1*) and Aiolos (*Ikzf3*). These Ikaros family factors are unique to B-lymphoid cells and function as transcriptional repressors and recruit components of the repressive nucleosome remodeling and histone-deacetylase (NuRD) complex^31-36^. Importantly, NuRD complex components (Chd4, Mta1, Mta2, Rbbp4, Gatad2a, Gatad2b, Mbd3, Hdac1, Hdac2) were identified as β-catenin-interacting proteins along with Ikzf1 and Ikzf3 (**Figure 4a-b**). To directly compare β-catenin-interacting protein complexes between human B-lymphoid, myeloid, and epithelial cells, we identified proteins bound to β-catenin in human B-ALL, B-cell lymphoma, myeloid, lung and colon cell lines by Co-IP and mass-spectrometry. Principal component analysis of β-catenin interactomes revealed that B-lymphoid cells (B-ALL, B-cell lymphoma) were clustered together along PC1 axis and separated from myeloid and colon and lung epithelial cells (**Figure 4c)**. In human B-lymphoid cells, IKZF1 and IKZF3 as well as the repressive NuRD components CHD4, RBBP4, MTA1, MTA2 and GATAD2B were among the most prominent interaction partners of β-catenin (**Figure 4d-e, S4a)**. In myeloid cells, β-catenin mainly interacted with a common core module of known interaction partners including CTNNA1, AXIN2, CTNNA2 and APC, that was also shared with all other cell types studied. In colon and lung epithelial cells, β-catenin preferentially interacted with TCF7L2 (**Figure 4e**) and histone acetyltransferases (KAT2B, TAF1; **Figure 4d, 4f**). Of particular interest is the epithelial cell-specific interaction of β-catenin with RUVBL1 (**Figure 4d, 4f**), which promotes β-catenin-mediated transcriptional activation of MYC^37-38^. Epithelial cell-specific binding of RUVBL1 to β-catenin was consistent with transcriptional activation of MYC by β-catenin-TCF7 complexes in epithelial cells^4-6^. In contrast, β-catenin induced repression of MYC in B-lymphoid cells and formed complexes with Ikaros factors and repressive NuRD components (**Figure 4d-f**).

**Figure 4:**
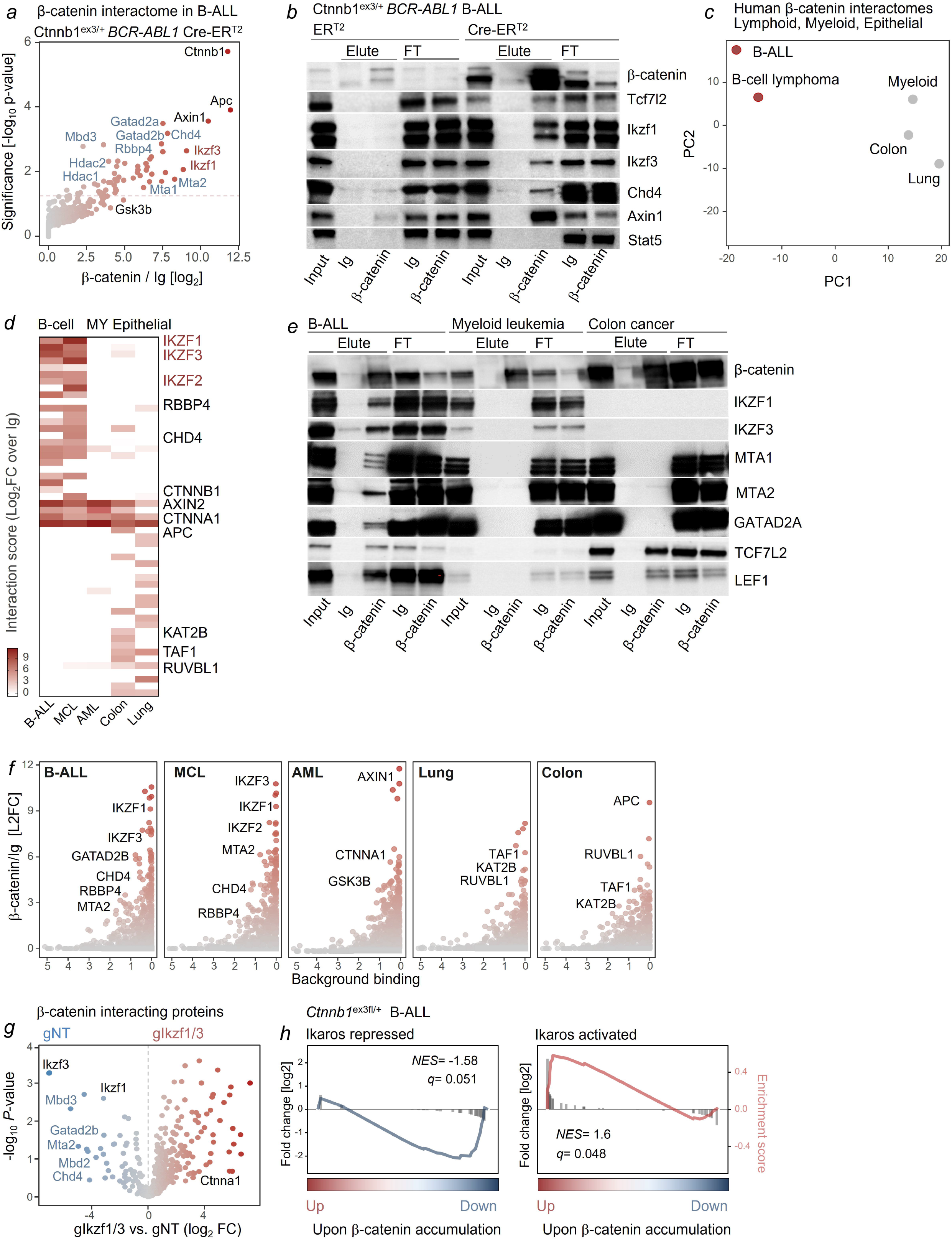
β-catenin forms repressive complexes with B-lymphoid transcription factors Ikzf1 and Ikzf3. (**a-b**) Proteins bound to β-catenin in B-ALL cells from *Ctnnb1*^ex3fl/+^ mice were enriched by co-IP, identified by mass spectrometry, and plotted based on statistical significance and log2-fold enrichment over IgG background control (n=4). Proteins with the most prominent binding to β-catenin included Ikaros factors Ikzf1 and Ikzf3 (red) and members of the repressive NuRD complex Chd4, Gatad2a, Gatad2b, Mta1, Mta2, Mdb3, Rbbp4, Hdac1, Hdac2 (blue). (**b**) β-catenin interacting proteins were validated by co-IP and Western blot in whole cell lysates (Input), proteins bound (Elute) and flow-through (FT) to isotype control or antibodies against β-catenin, using Stat5 as negative control. Co-IP experiments with antibodies against β-catenin or control Ig were performed in human B-ALL (MXP2), B-cell lymphoma (JEKO), AML (MOLM13), colon (SW480) and lung (H446) cancer cell lines expressing doxycycline inducible β-catenin. Eluted proteins were analyzed by mass-spectrometry. (**c**) Principal component analysis was performed to cluster cell lines based on similarity of β-catenin interactomes. (**d**) Heatmap of proteins that were enriched for β-catenin binding relative to Ig-control in B-ALL, mantle cell lymphoma (MCL), myeloid leukemia (AML), colon and lung cancer cell lines. (**e**) Whole cell lysates (Input), proteins bound and flow-through (FT) with β-catenin-antibodies or control Ig were analyzed by Western blotting to study interactions between β-catenin and Ikaros factors (IKZF1, IKZF3), NuRD complex components (MTA1, MTA2, GATAD2A) and TCF7L2, and LEF1 in B-ALL (PDX2), myeloid leukemia (JURL-MK1) and colon cancer (SW620) cells 16 hours following pharmacological β-catenin stabilization (LY2090314, 20 nM). (**f**) β-catenin binding proteins in each cell type were plotted as a function of background binding (x-axis, non-specific binding defined by CRAPOME database) and log2-fold enrichment over control Ig (y-axis). (**g**) Changes in β-catenin interactomes in B-ALL cells upon Ikaros factor deletion (gIkzf1/3) were analyzed by co-IP and mass-spectrometry. Proteins bound to β-catenin were plotted based on significance (y-axis) and log2-fold enrichment (x-axis) compared to B-ALL cells without deletion of Ikaros factors (gNT; n=3). (**h**) Amplification of Ikaros-mediated gene expression changes by β-catenin: depletion of genes repressed by Ikaros factors and enrichment of genes indirectly activated by Ikaros factors in murine B-ALL cells upon β-catenin accumulation.

### β-catenin-NuRD complex interactions depend on lymphoid-specific Ikaros factors

To test whether the unusual composition of the β-catenin interactome in B-lymphoid cells depends on lymphoid-specific Ikaros-factors, we deleted both Ikzf1 and Ikzf3 in murine B-ALL cells and repeated the Co-IP and mass spectrometry identification of β-catenin-interacting proteins. Deletion of *Ikzf1* and *Ikzf3* in B-ALL cells was achieved by electroporation-based delivery of Cas9 ribonucleoproteins (RNPs) containing Cas9 and guide-RNAs directed against *Ikzf1* (gIkzf1) and *Ikzf3* (gIkzf3) or a non-targeting control (gNT). Successful deletion was confirmed by Western blot screening of single clones for the loss of Ikzf1 and Ikzf3 (**Figure 5a**). Interestingly, deletion of Ikaros-factors resulted in loss of interactions between β-catenin and the NuRD complex components Chd4, Mbd3, Mbd2, Mta2 and Gatad2b (**Figures 4g**). Previous work demonstrated that loss of IKZF1 in B-ALL results in a shift from lymphoid to epithelial lineage features^32^. Consistent with these findings, our results suggest that β-catenin promotes, by default, transcriptional activation of an epithelial program, unless Ikaros factors redirect β-catenin to recruit repressive NuRD complexes characteristic of B-lymphoid cells.

**Figure 5:**
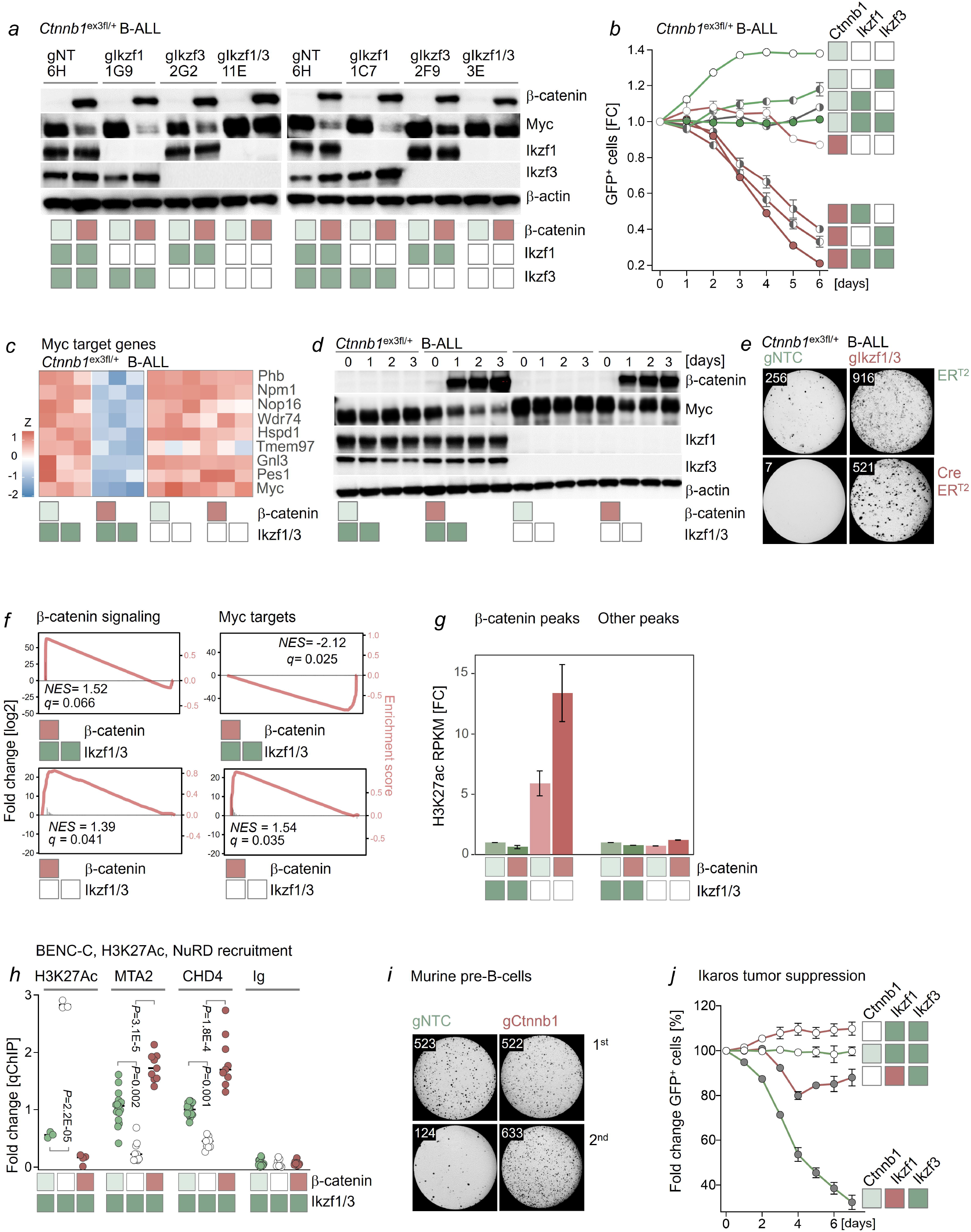
β-catenin functions as an amplifier of Ikaros-mediated gene expression changes. *BCR-ABL1*-transformed *Ctnnb1*^ex3fl/+^ B-ALL cells were gene-edited with crRNAs targeting Ikaros factors (*Ikzf1, Ikzf3*) individually or both or non-targeting crRNAs (gNT). Deletion of Ikaros factors was confirmed by Western blot in clonal cell lines established from single cells. Multiple clones were studied for each genotype. *Ctnnb1*^ex3fl/+^ B-ALL cells were transduced with 4-OHT-inducible GFP-tagged Cre-ER^T2^ or ER^T2^. Color code for boxes in **a-e**: light green box (β-catenin baseline), red box (β-catenin accumulated), dark green box (Ikaros factors baseline) and white box (Ikaros factors deleted). (**a**) Western blot was performed for β-catenin, Ikzf1, Ikzf3, Myc and β-actin two days after induction of Cre and β-catenin accumulation. (**b**) Competitive fitness of B-ALL clones was assessed in the presence or absence of β-catenin accumulation and deletion of either *Ikzf1, Ikzf3* or both Ikaros factors, using non-targeting crRNAs (gNT) as reference. (**c**) Heatmap to show changes in Myc target gene expression levels upon β-catenin activation with and without concurrent deletion of both Ikaros factors (Ikzf1, Ikzf3). (**d**) Western blot analyses to measure protein levels of β-catenin, Myc, Ikzf1 and Ikzf3 in relation to β-actin for 0-3 days after 4-OHT addition. (**e**) Colony forming assays for B-ALL cells with and without Ikaros factor deletion and with and without β-catenin accumulation (2 days) are shown. Representative images and colony numbers from three independent experiments are shown at 10 days after plating (triplicates). (**f**) GSEA plots for enrichment of β-catenin signaling (left) and MYC target genes (right) upon β-catenin accumulation and in the presence (bottom) or absence (top) of Ikaros factor deletion. (**g**) Quantification of changes in H3K27Ac ChIP-seq signals at β-catenin target regions vs. other regions following β-catenin accumulation in the presence or absence of Ikaros factor deletion. (**h**) ChIP-qPCR to measure enhancer activity (H3K27ac) and recruitment of NuRD complex components (MTA2 and CHD4) to the Myc superenhancer region (BENC-C) in B-ALL cells upon deletion of β-catenin (white circles) or accumulation of β-catenin (red circles) in comparison to wild type cells (green circles). Data were pooled from 7 independent qChIP experiments. (**i**) Murine B-ALL cells with and without engineered deletion of β-catenin were plated on methylcellulose. Primary (1^st^) and secondary (2^nd^) platings are shown, representative images and average counts of primary and secondary colonies from three independent experiments. (**j**) Murine B-ALL cells with (white box) and without engineered deletion of β-catenin (light green box) were transduced with vectors for inducible expression of GFP-tagged IKZF1 (red box) or GFP empty vector (green box). Changes in the frequencies of GFP^+^ cells were monitored by flow cytometry. Representative data from three independent experiments are shown.

### Ikaros-factors determine the outcome of β-catenin signaling

Since expression of Ikaros-factors and β-catenin are inversely correlated in B-lymphoid and epithelial cells (**Figure 1**), we tested whether ectopic expression of IKZF1 in epithelial tumor cell lines with constitutively high β-catenin protein levels caused transcriptional repression of MYC comparable to ectopic activation of β-catenin in Ikaros-expressing B-lymphoid cells. We transduced three colon and three lung cancer cell lines with constructs for doxycycline-inducible expression of IKZF1 or empty vector (EV) controls. Small molecule inhibition of GSK3β induced accumulation of β-catenin with slightly increased MYC expression in colon and lung cancer cell lines in the absence of IKZF1-expression. However, in the presence of ectopic IKZF1 expression, small molecule inhibition of GSK3β and accumulation of β-catenin had the opposite effect and suppressed MYC expression and induced cell death (**Figure S4b-c**). The outcome of β-catenin activation in colon and lung cells with ectopic IKZF1 expression was the same as in B-lymphoid cells with constitutive expression of Ikaros-factors. In a converse experiment, we determined whether the effects of β-catenin activation were dependent on B-cell identity and expression of B-lymphoid Ikaros-factors. Hence, we reprogrammed *Ctnnb1*^ex3fl/+^ B-ALL cells into the myeloid lineage by inducible expression of the myeloid transcription factor CEBPα. Two days after doxycycline-induced expression of CEBPα, B-ALL cells expressed the myeloid cell antigen CD11B (Mac1) and lost expression of CD19, Ikzf1 and Ikzf3 (**Figure S4d-e**). While Cre-mediated accumulation of β-catenin abolished competitive fitness and MYC expression in B-ALL cells, CEBPα-mediated myeloid-reprogramming fully restored clonal fitness and MYC expression levels (**Figure S4e-f**). Collectively, these findings suggest that Ikaros-factors determine the outcome of β-catenin signaling, namely transcriptional activation vs repression of MYC and other β-catenin/WNT targets. Beyond MYC, inducible accumulation of β-catenin broadly amplified Ikaros-mediated gene expression changes and deepened Ikaros-mediated repression and augmented transcriptional activation by Ikaros factors (**Figure 4h**).

### One single Ikaros factor is required and sufficient to enable β-catenin-induced repression of MYC

Deletion of *IKZF1* is a frequent lesion in B-ALL^39^, whereas *IKZF3* mutations are common in mature B-cell malignancies^40^. However, these lesions are typically monoallelic and cases with defects of both IKZF1 and IKZF3 are exceedingly rare. For this reason, we tested whether deletion of one single Ikaros factor, either *Ikzf1* or *Ikzf3* alone could rescue β-catenin-induced repression of MYC, survival and proliferation of B-ALL cells. While deletion of one Ikaros factor, either *Ikzf1* or *Ikzf3*, had no significant effects, only concurrent biallelic deletion of both B-lymphoid Ikaros factors reversed Myc-repression and cell death upon inducible accumulation of β-catenin (**Figure 5a-b**). This result suggests that the expression of one single Ikaros factor is required and sufficient for β-catenin-induced repression of MYC and induction of cell death. Indeed, in B-ALL cells harboring a biallelic deletion of either *Ikzf1* or *Ikzf3*, complex formation between β-catenin and the residual Ikaros factor remained intact, suggesting that heterodimerization between Ikzf1 and Ikzf3 is not required for complex formation with β-catenin (**Figure S5a**). In contrast, biallelic deletion of both *Ikzf1* and *Ikzf3* fully rescued colony formation, cell survival in a competitive cell culture assay and restored Myc-expression and Myc-driven transcriptional programs (**Figure 5b-e**). Measuring the effects of β-catenin-accumulation on enhancer activity (H3K27ac ChIP-seq) and gene expression (RNA-seq) in B-ALL cells, deletion of both Ikaros-factors largely erased effects of β-catenin-accumulation on gene expression, including repression of Myc (**Figures 5c, S5b-d**). These findings suggest that β-catenin activity is mainly directed by Ikaros factors and functions as an amplifier of Ikaros-dependent gene expression changes in B-lymphoid cells. Interestingly, β-catenin accumulation had opposite effects on Myc transcriptional programs, entirely depending on whether Ikaros factors were functional (repression) or deleted (activation; **Figure 5f**). In a genome-wide analysis, β-catenin accumulation suppressed enhancer activity and H3K27ac marks in the presence of functional Ikaros factors at β-catenin peaks (**Figure 5g**). Upon deletion of Ikaros factors, however, accumulation of β-catenin had the opposite effect and massively increased enhancer activity (H3K27ac) at β-catenin ChIP-seq peaks (**Figure 5g**). Together these findings suggest that β-catenin can, in principle, act as a powerful transcriptional activator in B-cells as in other cell types. However, B-lymphoid Ikaros factors reverse its positive effects on enhancer activity at β-catenin targets, including Myc.

Besides genetic ablation, we also studied pharmacological degradation of IKZF1 and IKZF3 by the cereblon modifier lenalidomide. Mechanistically, lenalidomide binds to the cereblon CRBN-CRL4 ubiquitin ligase to change its substrate affinity for selective ubiquitination and degradation of IKZF1 and IKZF3 proteins^41-42^. Here we show that lenalidomide not only induced efficient degradation of both IKZF1 and IKZF3 proteins in patient-derived B-ALL cells but also relieved β-catenin-induced transcriptional repression of MYC and suppression of colony formation (**Figure S6a-b**). Interestingly, lenalidomide treatment of patients with other hematological malignancies occasionally results in the development of B-ALL^43-44^. Hence, degradation of Ikaros-factors and derepression of MYC could be part of the underlying mechanism leading to the development-of lenalidomide-induced B-ALL in these cases.

### Ikaros factors redirect β-catenin from its canonical TCF7 binding sites

To determine how Ikaros factors and β-catenin interact at the chromatin level, we performed β-catenin, Ikzf1 and Ikzf3 ChIP-seq analyses to study changes of β-catenin- and Ikaros-binding peaks upon deletion of Ikaros factors or accumulation of β-catenin. Consistent with a dominant role of Ikaros factors in controlling β-catenin functions in B-cells, nearly 75% of β-catenin peaks were shared with Ikaros (Ikzf1, Ikzf3; **Figure S7a**). Inducible accumulation of β-catenin had very limited impact on Ikzf1 and Ikzf3 binding, ∼90% of Ikaros peaks remained unchanged (**Figure S7b**), with peaks in a Myc superenhancer region, termed blood enhancer cluster (BENC)^45^ among very few regions with increased Ikaros binding (**Figure S7c**). Likewise, β-catenin accumulation caused few changes in mRNA levels, including downregulation of Myc (**Figure S7d**). In contrast, loss of Ikaros-factors profoundly impacted β-catenin binding, affected about half of all β-catenin targets, generated 1,202 new β-catenin-binding peaks, while 1,675 β-catenin peaks were lost (**Figure S8a**). The Myc-BENC superenhancer region was among the regions with most prominent increases of β-catenin binding and increased Myc mRNA levels upon deletion of Ikaros factors (**Figure S7c-d**). When β-catenin accumulation was induced in the presence of Ikaros factors, new β-catenin peaks were mostly devoid of both H3K27ac and H3K4me3 marks. However, upon deletion of *Ikzf1* and *Ikzf3*, β-catenin peaks were substantially enriched for H3K27ac binding, suggesting increased enhancer activity at these sites. H3K4me3 marks were not changed upon Ikaros deletion (**Figure S8a**). The predominant increases of H3K27ac rather than H3K4me3 marks mirrored preferential interaction of β-catenin-Ikaros complexes with NuRD complex components, whereas histone methyltransferases and demethylases were not found among β-catenin-interacting proteins (**Figure 4d-g**). *De novo* β-catenin peaks associated with gain of H3K27ac marks suggest that deletion of *Ikzf1* and *Ikzf3* enabled redistribution of β-catenin to previously inactive enhancer regions to activate them. Motif enrichment analyses revealed that Ikaros-factor deletion restored targeting of β-catenin to classical TCF7, TCL7L1 and TCF7L2 motifs (**Figure S8a)**, consistent with canonical WNT signaling in epithelial cells^2-3^. These data suggest that Ikaros-factors sequester β-catenin away from transcriptional activation at canonical TCF7 sites. Consistent with a scenario in which Ikaros factors interfere with canonical β-catenin-TCF7 interactions in B-lymphoid cells, Ikaros factor deletion enabled or increased complex formation of β-catenin with TCF7-family transcription factors (Tcf7, Tcf7l1, Tcf7l2; **Figure S8b-c**).

### β-catenin is required for Ikaros-mediated tumor suppression

The B-lymphoid Ikaros family factors IKZF1 and IKZF3 function as important tumor suppressors in B-ALL^39^ and mature B-cell malignancies^40^, respectively. Unlike epithelial, myeloid, and T-cell lineage cells, B-lymphoid cells exhibit very low β-catenin protein expression and activity (**Figures 1, S1, S2**). Since β-catenin binds to NuRD complex components in an Ikaros-dependent manner and intensifies Ikaros-dependent gene expression changes (**Figure 4g-h**), we examined whether β-catenin, despite its low baseline expression levels, contributes to the recruitment of components of repressive NuRD complexes and regulation of enhancer activity at β-catenin/Ikaros target loci. To this end, we engineered genetic deletion of β-catenin in murine and human B-ALL cells and validated successful β-catenin deletion for single-cell clones by Western blot (**Figure S9a-b**). Genetic deletion of β-catenin did not significantly change enhancer activity (H3K27ac) at *Igll1, Myc* promoter and *Myc* epithelial enhancer regions. However, β-catenin-deletion markedly increased H3K27ac levels at BENC^45^ Myc superenhancer regions C and D (**Figures 5h, S9c**). Conversely, deletion of β-catenin modestly reduced recruitment of repressive NuRD complex components MTA2 and CHD4 at *Igll1, Myc* promoter and *Myc* epithelial enhancer regions, compared to near-complete loss of MTA2 and CHD4 recruitment at Myc superenhancer regions BENC-C and BENC-D (**Figures 5h, S9c**). Mirroring B-cell-specific functions of β-catenin-Ikaros complexes, loss of β-catenin significantly improved colony formation of B-lymphoid but not myeloid progenitor cells of human and murine origin (**Figures 5i, S9d-g**). Of note, this difference only became apparent in secondary replating experiments. Consistent with a role of β-catenin in negatively regulating B-cell leukemia-initiation in transplant experiments (**Figure 2f-g**), these results suggest that β-catenin-Ikaros complexes primarily limit self-renewal of B-lymphoid cells. To directly test a role of β-catenin in Ikaros-mediated tumor suppression, we compared the effect of inducible activation of IKZF1 in B-ALL cells with intact β-catenin (gNT) and B-ALL cells with genetic deletion of β-catenin (gCtnnb1). While IKZF1-induction induced cell death and suppressed proliferation in gNT B-ALL cells, deletion of β-catenin subverted IKZF1-mediated tumor suppression in gCtnnb1 B-ALL cells (**Figure 5j**). These results imply that despite low baseline expression levels, β-catenin, and its ability to engage Ikaros factors for the recruitment of repressive Ikaros:NuRD complexes represents a critical and previously unrecognized tumor suppressor in B-lymphoid malignancies.

### Both β-catenin and Ikaros factors are required for efficient NuRD complex recruitment

Deletion of Ikaros factors disrupted interactions between β-catenin and NuRD complex components (**Figure 4g**) and relieved β-catenin-induced repression of Myc (**Figures 5c-d, 5f**). Since β-catenin is required for effective NuRD complex recruitment (**Figure 5h, S9c**) and Ikaros-mediated tumor suppression (**Figure 5j**), we tested whether β-catenin and Ikaros factors cooperate in recruiting NuRD complex components. Consistent with cooperation between β-catenin and Ikaros factors, recruitment of the NuRD components MTA2 and CHD4 to BENC enhancer regions of Myc was increased by accumulation of β-catenin, but nearly entirely lost upon deletion of Ikaros factors or deletion of β-catenin (**Figure 6a, Figure S10**). In addition to BENC *Myc* enhancer regions, deletion of Ikaros and β-catenin also affected NuRD complex recruitment at the *Myc* promoter and the Ikaros target gene *Igll1* but not epithelial *Myc* enhancer regions (**Figure S10a**). Collectively, these results suggest that both Ikaros and β-catenin are required for effective NuRD complex recruitment and transcriptional repression of Myc in B-lymphoid cells.

**Figure 6:**
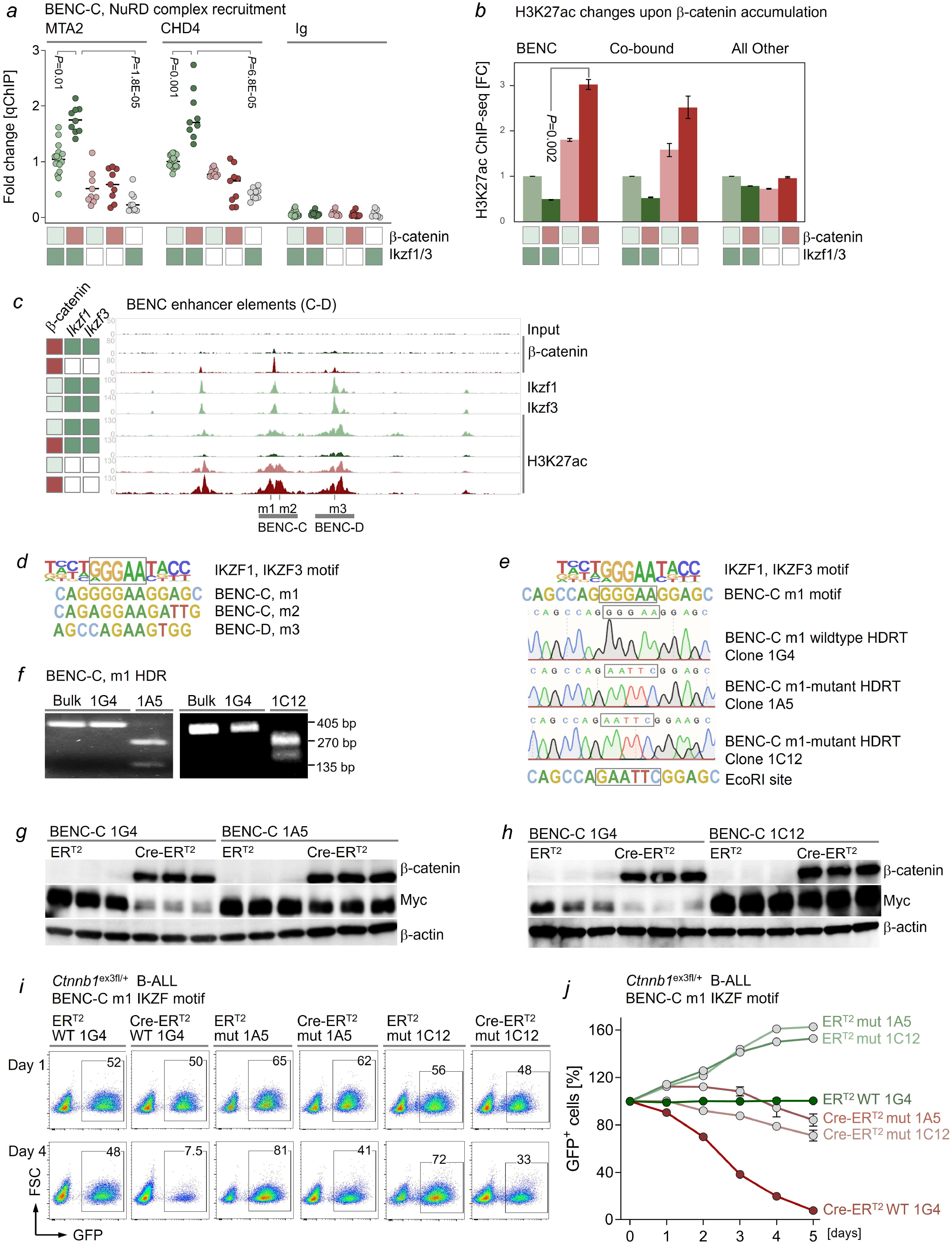
Mutation of a single Ikaros-motif of the BENC-C region subverts β-catenin-mediated repression of MYC. (**a**) ChIP-qPCR analysis of recruitment of NuRD complex components (MTA2 and CHD4) to the BENC-C enhancer region in *Ctnnb1*^ex3fl/+^ B-ALL cells. Genotypes are denoted by light green boxes (β-catenin baseline), red boxes (β-catenin accumulation), dark green boxes (Ikaros factors baseline) and empty boxes (Ikaros-null or β-catenin-null). Data represent a pool of 6 independent experiments. (**b**) Quantification of H3K27ac ChIP-seq signals at BENC enhancer regions, other regions with binding of both Ikaros factors and β-catenin (Co-bound) and all other regions. H3K27ac ChIP was performed with and without β-catenin accumulation and in the presence or absence of Ikaros factor deletion. (**c**) BENC elements C and D were analyzed for changes in β-catenin, Ikzf1, and Ikzf3 binding and H3K27 acetylation, upon induction of β-catenin in B-ALL cells with and without deletion of Ikaros factors. (**d**) Identification of Ikaros binding motifs in the BENC-C (m1, m2) and BENC-D (m3) elements. (**e-j**) Homology directed repair (HDR)-mediated editing of the BENC-C m1 motif to generate a new *Eco*RI site. To abrogate the binding of Ikzf1 and Ikzf3, the Ikaros core motif GGGAA was mutated, and clonal cell lines were generated and analyzed by (**e**) Sanger sequencing and (**f**) *Eco*RI digestion and gel electroporation. (**g-h**) Western blot analysis of Myc protein levels one day after β-catenin accumulation in B-ALL cells carrying intact or mutated BENC-C Ikaros m1 motifs. (**i-j**) Growth kinetics of B-ALL cells with intact and mutant BENC-C Ikaros m1 motif following Cre-mediated induction of β-catenin. (**i**) Representative FACS plots and (**j**) changes in the percentages of GFP^+^ cells are depicted.

### IKZF1 and IKZF3 mediate transcriptional repression of MYC at BENC superenhancer regions

Studying H3K27ac signals across multiple *MYC* super-enhancer clusters revealed that most of the enhancer activity was concentrated in blood enhancer cluster (BENC) regions (**Figure S10b**), which was identified as critical for the regulation of MYC expression in B-lymphoid and other hematopoietic cells^45^. Consistent with predominant recruitment of NuRD complex components at these regions (**Figures 6a, S10a**), Ikaros factors and β-catenin strongly bound to elements C-D of the BENC region (**Figures 6b-c, S10b**). In the presence of Ikaros factors, β-catenin-accumulation suppressed H3K27ac signals at BENC regions. However, in the absence of Ikaros factors, β-catenin-accumulation had the opposite effect and significantly increased H3K27ac signals at BENC enhancer regions (**Figures 6b-c, S10b**). Interestingly, other loci that were bound by both Ikaros factors and β-catenin showed a similar pattern (**Figure 6b**). While β-catenin-Ikaros complexes suppressed MYC-expression in B-lymphoid cells, these observations suggest that deletion of *Ikzf1* and *Ikzf3* releases β-catenin from transcriptional repression and restores its ability to promote transcriptional activation of MYC as in non-lymphoid cell types^4-6^.

### Mechanistic basis of β-catenin-mediated repression of MYC

Sequence analysis of BENC-C and BENC-D regions identified three Ikaros binding motifs (m1-m3) at significant Ikzf1 and Ikzf3 ChIP-seq peaks (**Figure 6c-d**). Of these, m1 perfectly matched the Ikaros motif (GGGAA), whereas the other two had a single base pair mismatch. To test the functional significance of the Ikaros m1 motif within the *Myc* BENC-C superenhancer region, we engineered knockin alleles to replace the Ikaros binding motif with an *Eco*RI site. After HDRT-based knockin of wildtype and mutant BENC-C alleles into murine *Ctnnb1*^ex3fl/+^ B-ALL cells, clones carrying the knockin mutation were selected based on *Eco*RI digestion and confirmed by Sanger sequencing (**Figure 6e-f**). *Ctnnb1*^ex3fl/+^ B-ALL clones with wildtype and mutant BENC-C Ikaros motifs were transduced with inducible Cre for accumulation of β-catenin. As expected, BENC-C wildtype knockin clones rapidly lost Myc expression and underwent cell death upon inducible accumulation of β-catenin (**Figure 6g-j**). In contrast, *Ctnnb1*^ex3fl/+^ B-ALL clones carrying knockin alleles for the mutant Ikaros m1 motif in BENC-C expressed Myc at higher baseline levels and were resistant to inducible β-catenin accumulation. Upon β-catenin accumulation, Myc levels remained high. Cell viability and competitive fitness of B-ALL clones carrying the mutant Ikaros m1 motif remained largely unchanged (**Figure 6g-j**). While it is likely that β-catenin accumulation has other effects in B-ALL cells, these findings underscore that β-catenin-induced toxicity and repression of Myc primarily depend on its interactions with Ikaros factors and in particular one single Ikaros binding site within the *Myc* BENC-C enhancer region.

### Pharmacological engagement of β-catenin-Ikaros complexes for refractory B-cell malignancies

Given that low baseline expression levels of β-catenin were sufficient to enable tumor suppression by Ikaros-factors in B-ALL cells (**Figure 5j**), pharmacological accumulation of β-catenin could potentiate tumor suppressive effects of β-catenin-Ikaros complexes. Our genetic approaches achieved β-catenin accumulation based on Cre-mediated excision of GSK3β-phosphorylation sites^30^. This previously unrecognized strategy to engage β-catenin-Ikaros complexes would be orthogonal to conventional mechanisms of drug-resistance and potentially useful in the treatment of patients with relapsed or refractory B-cell malignancies. For this reason, we next tested pharmacological approaches of β-catenin accumulation based on small molecule inhibitors of GSK3β. To address potential safety concerns related to pharmacological β-catenin accumulation, we focused our analysis on compounds that have completed clinical development and demonstrated favorable safety profiles in clinical trials (**Figure S11**).

For proof-of-concept studies, we tested six small molecule GSK3β inhibitors for their ability to selectively kill B-lymphoid leukemia and lymphoma cells. Four of the six GSK3β inhibitors (LY2090314, 6-Bromoindirubin-3’-oxime, CHIR98014 and CHIR99021) induced cell death at low nanomolar concentrations selectively in B-lymphoid but not myeloid and epithelial cells (**Figure S11a-b**). In contrast, Tideglusib had no significant activity in any cell type, while 9-ING-41 showed broad non-specific toxicity across all cell types tested (**Figure S11a-b**). Interestingly, the four GSK3β inhibitors with B-cell-selective toxicity (LY2090314, 6-Bromoindirubin-3’-oxime, CHIR98014 and CHIR99021) induced massive accumulation of β-catenin in parallel with acute suppression of MYC (**Figure 7a, 7c, S11c**). In contrast, lack of specific drug-responses for 9-ING-41 and Tideglusib was mirrored by failure to induce β-catenin accumulation and MYC-suppression (**Figure S11c**). These results suggest that accumulation of β-catenin and MYC-suppression not only represent important biomarkers for drug-responses to GSK3β-inhibitors but also reflect their underlying mechanism of action.

**Figure 7:**
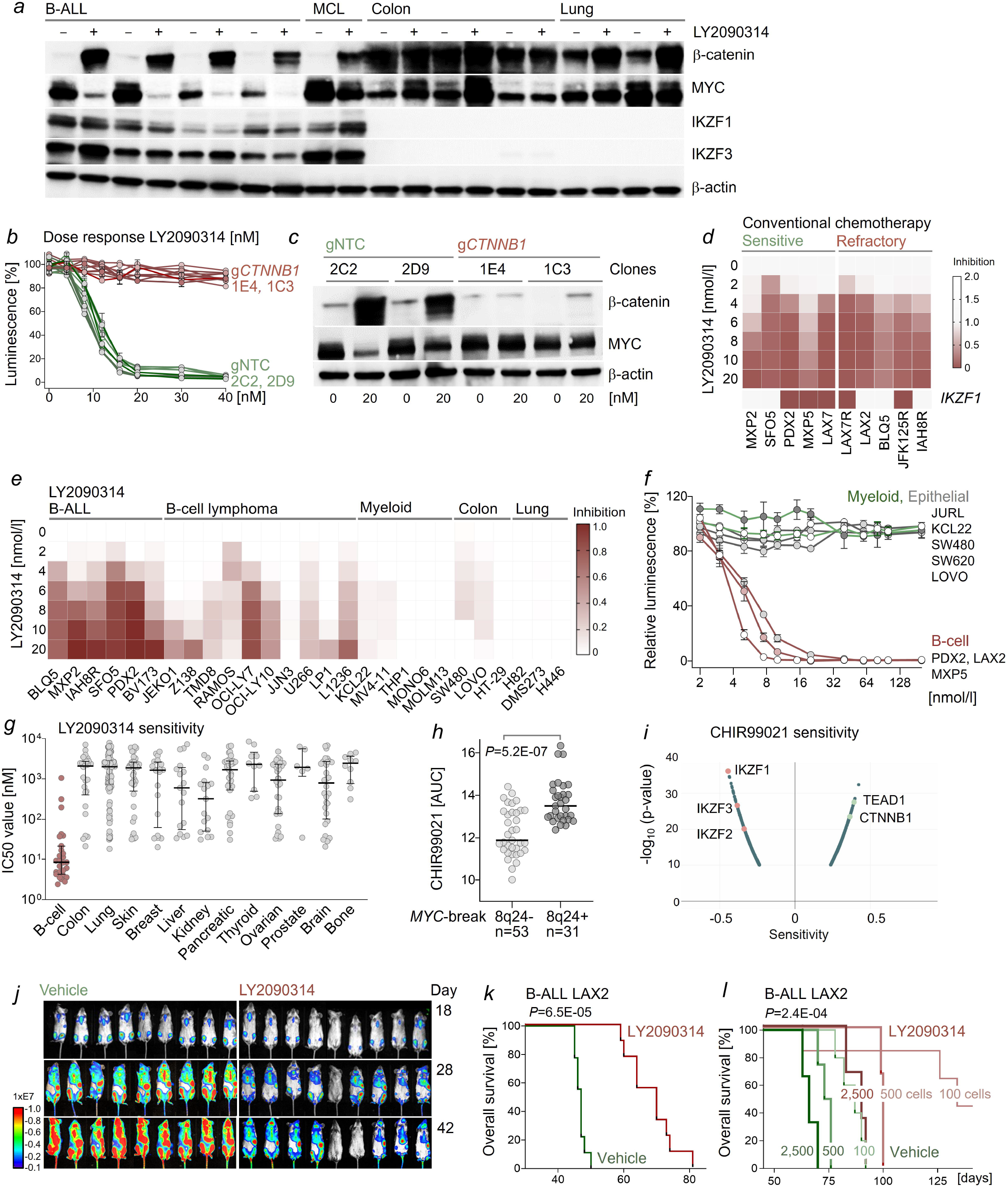
Pharmacological engagement of β-catenin-Ikaros complexes for targeted repression of MYC. (**a**) B-ALL (MXP2, LAX2, BLQ5, IAH8R), mantle cell lymphoma (MCL; JEKO1), colon (SW480, LOVO, HT-29), and lung cancer (H82, H446) cell lines were treated with the GSK3β small molecule inhibitor LY2090314 (20 nM) for one day. β-catenin, MYC and IKZF1 protein levels were assessed by Western blot, using β-actin as loading control. (**b**) Human B-ALL cells (BV173) were edited with crRNAs targeting β-catenin (g*CTNNB1*) or non-targeting crRNAs (gNT) and single cell-derived colonies were generated (**Figure S9b**). B-ALL cells with (clones 1E4, 1C3; g*CTNNB1*) or without (clones 2C2, 2D9; gNT) deletion of β-catenin were treated with LY2090314 for 3 days at the indicated concentrations and relative viability was determined by luminescence measurements. (**c**) Human B-ALL cells (BV173) with and without deletion of *CTNNB1* were treated with LY2090314 (20 nM) for 16 hours to force accumulation of β-catenin. Western blot was performed to analyze β-catenin and MYC levels. (**d**) Growth inhibition by the GSK3β inhibitor LY2090314 was compared for human B-ALL samples from patients who responded to conventional chemotherapy (sensitive) and from patients with refractory B-ALL (refractory). (**e**) Sensitivity to LY2090314 was assessed in a panel of 28 B-ALL, B-cell lymphoma, myeloid leukemia, colon, and lung cancer cell lines. Growth inhibitory effects were shown as heatmap. (**f**) B-ALL cells (red), myeloid leukemia (green) and colon cancer (gray) cell lines were treated with LY2090314 at concentrations between 0 up to 200 nM for 3 days and cell viability was determined by normalizing the luminescence signal of treated cells to untreated cells. (**g**) Responses to LY2090314 in 343 epithelial cancer cell lines (Prism Drug Repurposing Secondary Screen)^46^ and 17 B-lymphoid cell lines (B-ALL, 7 B-cell lymphoma; red circles) were plotted as IC_50_ values (nM). (**h**) Drug responses (AUC) to the GSK3β-inhibitor CHIR99021 were plotted for 84 B-cell lymphomas with (8q24+; n=31) and without (8q24-; n=53) *MYC* rearrangement. (**l**) Computational analyses of gene expression (biomarker) correlations with responses to the GSK3β-inhibitor CHIR99021 in epithelial cancers and B-lymphoid cell lines^46^. Expression of Ikaros-factors was positively associated with sensitivity to CHIR99021, while expression of β-catenin and the epithelial marker TEAD1 correlate with CHIR99021-resistance. (**j-l**) Luciferase-labelled LAX2 cells were injected into sub-lethally irradiated NSG mice. Mice were either treated with 10 mg/kg LY2090314 or vehicle control. (**j**) Leukemia burden was assessed by bioluminescence imaging at day 18 (*top*), 28 (*middle*) and 42 (*bottom*) following transplantation. (**k**) Kaplan-Meier analysis of overall survival in each group (n=9, *P*=6.5E-05; calculated by Logrank test). (**l**) Effect of LY2090314 on leukemia-initiation was studied by transplanting limiting doses (100-2,500 cells) of B-ALL cells prior to treatment into sub-lethally irradiated NSG mice.

### β-catenin accumulation represents the mechanism of action of GSK3β-inhibitors in B-cell malignancies

To determine if accumulation of β-catenin indeed represents the mechanistic basis of LY2090314-mediated cell death in B-ALL cells, we deleted *CTNNB1* in human B-ALL cells using Cas9-RNPs and screening of clones for *CTNNB1*-deletion from single cells (**Figure S9b**). Reminiscent of knockin mutation of the m1 Ikaros binding motif within the *Myc* BENC-C superenhancer region (**Figure 6g-j**), deletion of *CTNNB1* conferred near-complete resistance of B-ALL cells to LY2090314 and prevented suppression of MYC (**Figure 7b-c**). In our B-ALL mouse model, we had shown that one single Ikaros factor (IKZF1 or IKZF3) was sufficient to suppress MYC and induce cell death upon β-catenin accumulation (**Figure 5a-b**). Since *IKZF1*-deletions are common in human B-ALL^39^, we tested the impact of *IKZF1*-deletion on sensitivity to the GSK3β inhibitor LY2090314. Studying 10 patient-derived xenografts (PDX), including 5 with *IKZF1*-deletion, we found no significant differences in responses to LY2090314 (**Figure 7d**). In addition, whether or not B-ALL PDX were derived from patients who responded to standard chemotherapy or were refractory and relapsed, did not affect responses to LY2090314. This result suggests that pharmacological β-catenin accumulation by GSK3β-inhibition is indeed orthogonal to conventional mechanisms of drug-resistance and may represent a vulnerability that could be impactful for patients with drug-resistant or relapsed B-cell malignancies. Focused analyses of drug-responses for LY2090314 in a larger panel of cell lines and PDX corroborated profound responses in B-ALL and B-cell lymphoma cells in the absence of significant effects on myeloid and epithelial tumor cells (**Figure 7e-f**). A combined analysis of responses to LY2090314 based on 343 epithelial cancer cell lines (Prism Drug Repurposing Secondary Screen)^46^ and B-lymphoid cell lines (17 B-ALL, 7 B-cell lymphoma) revealed that IC_50_ values for LY2090314 were substantially higher in epithelial and myeloid cancer cells than in B-ALL (236-fold) and B-cell lymphoma (92-fold; **Figure 7g**). While B-ALL cell lines were uniformly sensitive, sensitivity to the GSK3β-inhibitor CHIR99021 showed a bimodal distribution across 84 B-cell lymphoma cell lines. The difference between the two groups largely tracked with presence or absence of *MYC* translocations: B-cell lymphoma cell lines with *MYC*-rearrangement (n=31; 8q24+) were substantially less sensitive to GSK3β-inhibition compared to cell lines without *MYC*-translocation (n=53; *P*=5.2E-07; **Figure 7h**). Computational analyses of additional biomarkers for responses to GSK3β-inhibition across Prism panel of cell lines identified Ikaros factor (IKZF1, IKZF3, IKZF2) expression as the top-ranking association, while high baseline levels of β-catenin and the epithelial cell transcriptional factor TEAD1 showed the strongest negative correlation with sensitivity to CHIR99021 (**Figure 7i**).

### Rationale for repurposing of GSK3β inhibitors for refractory B-lymphoid malignancies

Five GSK3β-inhibitors, including LY2090314 and CHIR99021, had previously achieved favorable safety and PK/PD profiles at micromolar plasma concentrations (Cmax) in 22 clinical trials for patients with pancreatic cancer, advanced sarcoma, Alzheimer’s disease, progressive supranuclear palsy, amyotrophic lateral sclerosis, tooth repair, hearing loss and NK-cell stimulation for immunotherapy (**Figure S11d**). Since LY2090314 and CHIR99021 demonstrated B-cell selective activity at low nanomolar concentrations (**Figures 7e-f, S11b**), accumulation of β-catenin and suppression of MYC (**Figures 7a, S11c**), we tested a potential rationale for repurposing GSK3β small molecule inhibition towards refractory B-cell malignancies. To this end, we treated immunodeficient mice, bearing patient-derived xenografts (PDX) from refractory B-ALL cells with LY2090314 as a single agent. Sublethally irradiated (2 Gy) NSG mice bearing refractory B-ALL PDX were injected intraperitoneally with LY2090314 or vehicle control for six times. Compared to mice treated with vehicle, LY2090314 substantially reduced leukemia burden and significantly extended overall survival (*P*=6.5E-05, n=9; **Figure 7j-k**). Consistent with highly B-cell-selective effects of GSK3β-inhibition, treated mice did not show significant weight loss or other dose-limiting toxicity. These findings suggest that GSK3β-inhibition, when used in combination with existing regimen for refractory B-ALL and other lymphoid malignancies, could substantially deepen treatment responses and overcome mechanisms of conventional drug resistance. GSK3β small molecule inhibitors, including LY2090314, AZD1080, Laduviglusib (CHIR99021), Tideglusib and Elraglusib (9-ING-41), have demonstrated safety and tolerability at micromolar plasma concentrations (Cmax)^47-53^. Among the reported adverse effects of LY2090314 in clinical trials (Cmax micromolar) was lymphopenia (**Figure S11d**), which is consistent with the unique dependency of B-lymphoid cells on GSK3β-mediated degradation of β-catenin discovered in this study. Importantly, our studies of LY2090314 and Laduviglusib (CHIR99021) in refractory B-cell malignancies demonstrated B-cell selective effects at low nanomolar concentrations *in vitro* (**Figures 7b, 7d-g, S11b-c**) and *in vivo* (**Figures 7j-l**). Consistent with impairment of B-cell leukemia-initiation upon genetic accumulation of β-catenin (**Figure 2f-g**), pharmacological β-catenin accumulation by LY2090314 reduced the leukemia-initiation potential of B-ALL PDX in transplant recipient mice (**Figure 7l**).

## DISCUSSION

Interestingly, a recent forward genetic screen for mutations affecting lymphopoiesis in mice discovered *Lmbr1l* as an essential negative regulator of β-catenin^54^. Loss of *Lmbr1l* resulted in profound B- and T-cell defects. Consistent with our finding of GSK3β-mediated degradation of β-catenin as unique B-lymphoid vulnerability, Lmbr1l is required to stabilize GSK3β and lymphopoiesis in *Lmbr1l*-deficient mice could be rescued by deletion of β-catenin^54^. Consistent with these findings, β-catenin deletion in our study rescued the toxic effect of GSK3β small molecule inhibition in B-lymphoid cells (**Figure 7b-c**).

### β-catenin-Ikaros complexes in T-cells

Of note, loss of *Lmbr1l* resulted in β-catenin accumulation and profound defects of both B- and T-lymphopoiesis. This would be consistent with expression and activity of some Ikaros factors (e.g. IKZF1) in both B- and T-lymphoid cells^55^. However, unlike B-lymphoid cells, T-cells exhibit substantial baseline activity of β-catenin signaling (**Figure 1c, S1**) and T-cell malignancies carried activating β-catenin mutations at similar frequencies as in solid tumors (**Table S1**). In some T-cell malignancies, oncogenic activation of Notch1 counteracts Ikaros-mediated tumor suppression^56^, which could provide a mechanism for T-lymphoid cells to become permissive to β-catenin accumulation. Seemingly contrasting our scenario that β-catenin-Ikaros complexes suppresses lymphoid development, targeted overexpression of β-catenin in thymocytes resulted in the development of T-lymphoid malignancies^25, 57^. Strikingly, karyotypic analyses of 18 β-catenin-driven T-cell lymphomas in two studies revealed that 17 of them carried a *Myc*-rearrangement^25, 57^. While Myc is a target of transcriptional activation by β-catenin in other cell types^4-6^, these results suggest that β-catenin itself did not promote Myc expression in T-cells and instead imposed selective pressure for secondary genetic lesions resulting in Myc-overexpression: In the first study, all 8 β-catenin-driven T-cell lymphomas harbored a *Myc* translocation, including *Myc-Tcra* (n=6) and *Myc-Tcrb* (n=1) rearrangements^25^. In the second study, 9 of 10 β-catenin-driven T-cell lymphomas carried either a *Myc-Tcra* (n=3) rearrangement or large deletions downstream of *Myc* encompassing the BENC region (n=6)^57^. The remaining case without *Myc* abnormality showed aberrant overexpression of N-Myc^57^. Consistent with our finding that targeted mutation of an Ikaros motif in the Myc BENC-C region conferred resistance to β-catenin accumulation (**Figure 6**) and that *MYC*-translocations in B-cell lymphomas are correlated with reduced sensitivity to GSK3β-inhibition (**Figure 7h**), these findings in T-cell malignancies^25, 57^ raise the possibility that β-catenin may form similar complexes with Ikaros factors for repression of *Myc* in T-cells. Studies to explore β-catenin-Ikaros complexes and the role of Myc BENC enhancer regions in normal T-cell development and T-cell malignancies are currently underway.

### Limitations of our proposal to repurpose GSK3β-inhibitors for refractory B-cell malignancies

In agreement with an early study suggesting that Lef1/β-catenin signaling may negatively regulate Myc expression in B-cells^58^ -as opposed to other cell types^4-6^, we propose here to leverage clinically approved GSK3β-inhibitors to engage β-catenin-Ikaros complexes for targeted repression of Myc as a previously unrecognized strategy to overcome drug resistance in refractory B-cell malignancies.

Translocations of the *MYC* gene at 8q24 occur in about 15% of all B-cell malignancies^59-60^. In many of these cases, expression of translocated *MYC* is driven by the *IGH* Eµ enhancer and *IGH* 3’ regulatory regions and no longer regulated by its transcriptional control elements (e.g., BENC-C). Our experiments based on CRISPR-based knockin mutation of a single Ikaros-binding motif identified the lymphoid *Myc* BENC-C superenhancer region as a central mechanistic element in β-catenin-dependent Myc repression (**Figure 6**). On this basis, we predict that GSK3β-inhibition in B-cell lymphomas with *MYC* translocation will likely fail to repress MYC. Consistent with this scenario, B-cell lymphoma cell lines carrying a *MYC*-translocation at 8q24 were substantially less sensitive to GSK3β-inhibition (**Figure 7h**). These findings are in line with a previous study of a genetic mouse B-cell lymphoma model for overexpression of MYC from the *IGH* Eµ-enhancer, which enabled secondary lesions resulting in β-catenin-hyperactivation^61^. In our survey of 2,137 B-cell malignancies, we found 17 cases with an activating β-catenin pathway lesion (**Figure 1b, Table S1**). Among 1,980 B-cell lymphomas with informative MYC-status and without β-catenin pathway lesion, 208 carried a *MYC* break at 8q24 (11%). Reminiscent of mouse models with T-cell-specific overexpression of β-catenin^25, 57^, among 14 cases with β-catenin pathway lesions and informative *MYC*-status, 12 carried a *MYC*-break (expected 1.5, observed 12, χ² test *P*=3.6 E-5; **Table S2)**.

Based on genetic knockin mutation of a single Ikaros-binding motif we demonstrated the critical importance of the Myc BENC-C superenhancer region as mechanistic basis for GSK3β-inhibitor activity (**Figure 6**). Recent work demonstrated far-reaching effects of aberrant somatic hypermutation targeting superenhancer regions, including *MYC*, in diffuse large B-cell lymphomas (DLBCL)^62^. Studying *MYC* BENC superenhancer regions in 93 DLBCL cases and normal germinal center B-cells as a reference, 89 point mutations were found in 49 cases (52.6%, range 1-8 mutations per case) as byproduct of aberrant somatic hypermutation (**Table S3**). While the Ikaros m1 motif within MYC BENC-C (**Figure 6**) was not mutated, these results show that the BENC region is subject to pervasive hypermutation in DLBCLs and sporadic mutations of the Ikaros motif within the BENC-C region could be a mechanism to confer resistance to GSK3β-inhibition as observed in our CRISPR experiment with an engineered knockin mutation in the Ikaros m1 motif within BENC-C (**Figure 6**). In addition, deletion of MYC downstream regions, including BENC, and aberrant overexpression of N-Myc -as observed in T-cell lymphomas^57^, could represent mechanisms that confer resistance to GSK3β-inhibitor treatment.

B-ALL, unmutated CLL and mantle cell lymphomas are derived from pre-germinal center stages of B-cell development that are not subject to somatic hypermutation. In addition, *MYC*-rearrangements are exceedingly rare in B-ALL, unmutated CLL and MCL. In our experiments, B-ALL and mantle cell lymphomas were highly sensitive to GSK3β-inhibition *in vitro*, suggesting that patients with these diseases might benefit from a targeted repurposing effort of GSK3β-inhibitors.

For the past 60 years, glucocorticoids have been central empirical components of nearly all treatment regimen for B-lymphoid malignancies^63^. Likewise, L-asparaginase and methotrexate^64-65^ have highly selective effects on B-lymphoid malignancies with very limited toxicity in myeloid and epithelial cells^63-65^. More recently, antibody-mediated killing (e.g., rituximab) and inhibition of B-cell receptor signaling (e.g. ibrutinib^66-67^) have built on the concept of targeted elimination of B-lymphoid cells, with B-cell depletion and lymphopenia as acceptable side-effect. While these treatments have revolutionized the treatment of B-cell malignancies, the frequent development of resistance (e.g. to glucocorticoids in B-ALL and ibrutinib in B-cell lymphomas) has limited their overall success. Given their B-cell-selective activity, our future repurposing efforts of GSK3β-inhibitors will focus on opportunities to overcome resistance to glucocorticoids in B-ALL and ibrutinib in B-cell lymphomas.

## Acknowledgments

We would like to thank Dr. Laura Pasqualucci and Dr. Riccardo Dalla-Favera for critical discussion and MYC enhancer mutation data, Lars Klemm and current and former members of the Müschen laboratory for their support and helpful discussions. Research in the Müschen laboratory is supported by the NIH through an NCI Outstanding Investigator Award R35CA197628, R01CA157644, R01CA213138, R01AI164692 and P01CA233412 (to M.M.), the Howard Hughes Medical Institute HHMI-55108547 (to M.M.), the Arthur H. and Isabel Bunker Chair in Hematology (to M.M.), a Blood Cancer Discoveries Grant program through The Leukemia & Lymphoma Society, The Mark Foundation for Cancer Research, and The Paul G. Allen Frontiers Group and the V Foundation for Cancer Research T2018-003B (to M.M.). M.M. is a Howard Hughes Medical Institute (HHMI) Faculty Scholar.

The authors have no financial conflicts of interest.

