## Supplementary Figures for "Targeted engagement of β-catenin-Ikaros complexes in refractory B-cell malignancies"

**Figure S1:** *Lack of β-catenin expression and activity in B-lymphoid cells*

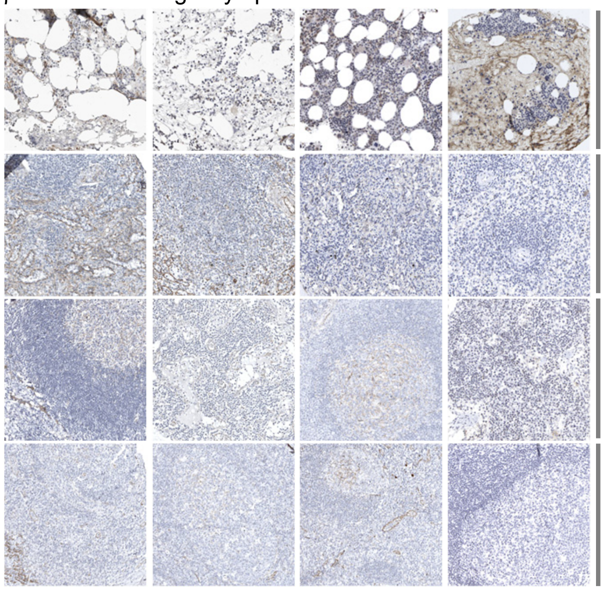


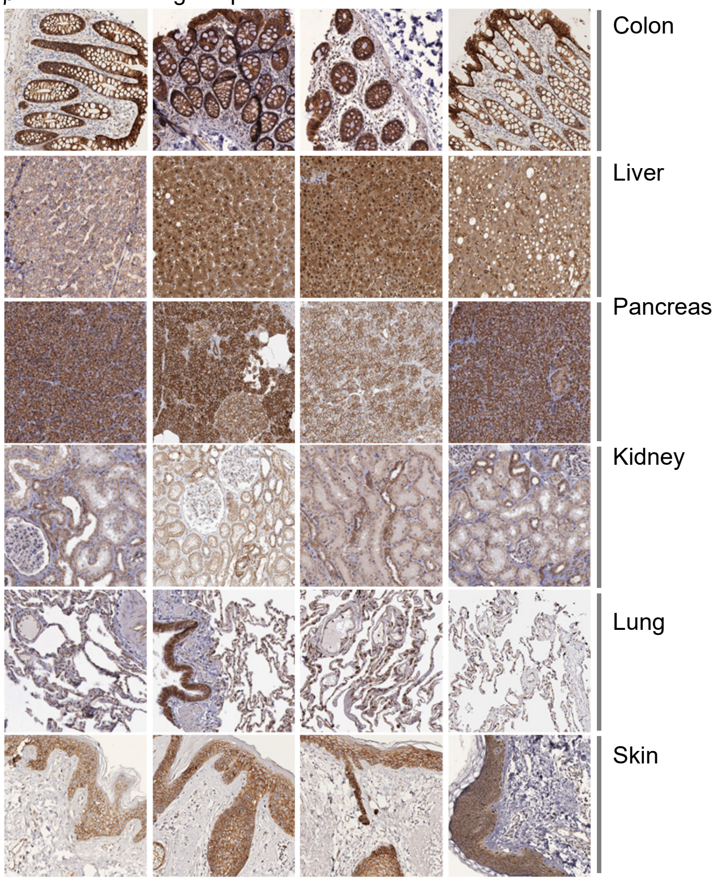


(**a**) Immunohistochemical staining for β-catenin expression (brown, H&E counterstain) in human lymphoid tissues (*left*), including bone marrow (n=7), spleen (n=6), lymph node (n=8), tonsil (n=9) as well as epithelial tissues (*right*), including colon (n=7), liver (n=9), pancreas (n=12), kidney (n=8), lung (n=8) and skin (n=7). Representative images are shown.

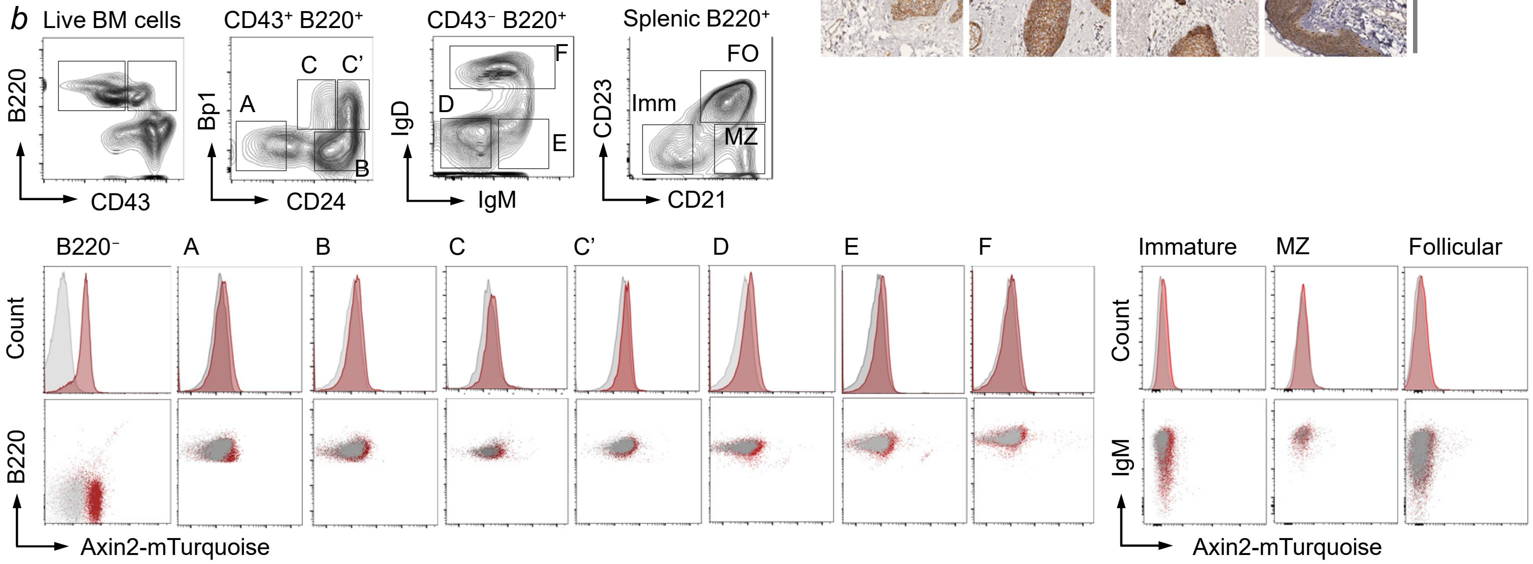


β-catenin signaling was measured in B-lymphoid (**b**) and T-lymphoid (**c**) cells in mice carrying the Axin2-mTurquoise transgene[28] (red) relative to background signal in control mice lacking the reporter transgene (gray). Data shown are representative of three mice from two independent experiments. (**b**) For B-lymphoid cells, bone marrow B220$^+$ CD43$^+$ B-cell progenitors were separated as Fraction A (Bp1$^-$ CD24$^-$), B (Bp1$^-$ CD24$^+$), C (Bp1$^{lo}$ CD24$^+$), C' (Bp1$^{hi}$ CD24$^+$) and B220$^+$ CD43$^-$ B-cells were classified as Fraction D (IgM$^-$ IgD$^-$), E (IgM$^+$ IgD$^-$) and F (IgM$^+$ IgD$^+$). Among B220$^+$ splenic B-cells, immature (Immature, CD21$^-$ CD23$^-$), marginal zone (MZ, CD21$^+$ CD23$^-$) and follicular (CD21$^+$ CD23$^+$) B-cells were studied. (**c**) β-catenin signaling was measured in CD4$^-$ CD8$^-$ double negative (DN), CD4$^+$ CD8$^+$ double positive thymocytes as well as CD4$^+$ and CD8$^+$ single positive T-cells. DN thymocytes were further separated as DN1-4 based on CD25 and CD44 expression.

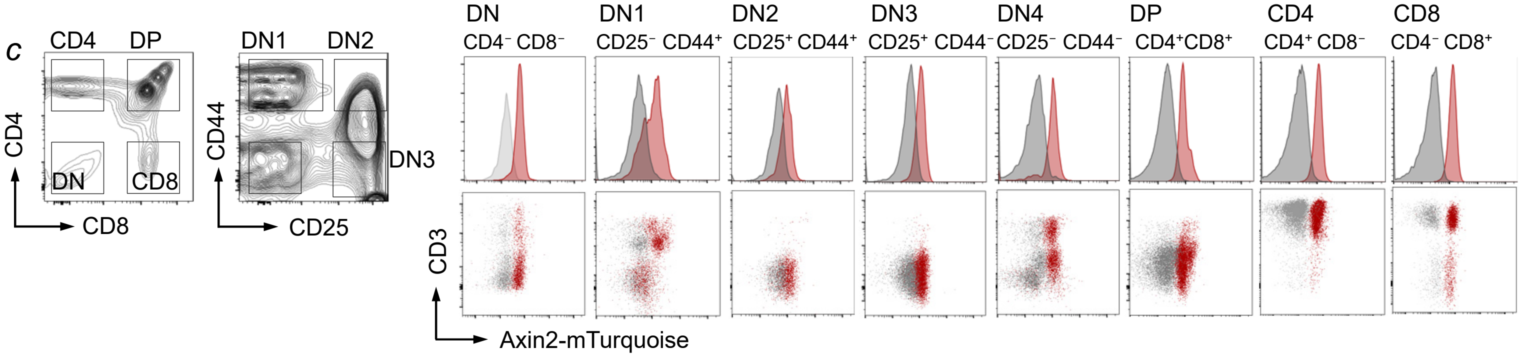

**Figure S2:** *Lack of β-catenin expression and activity in B-cell malignancies*

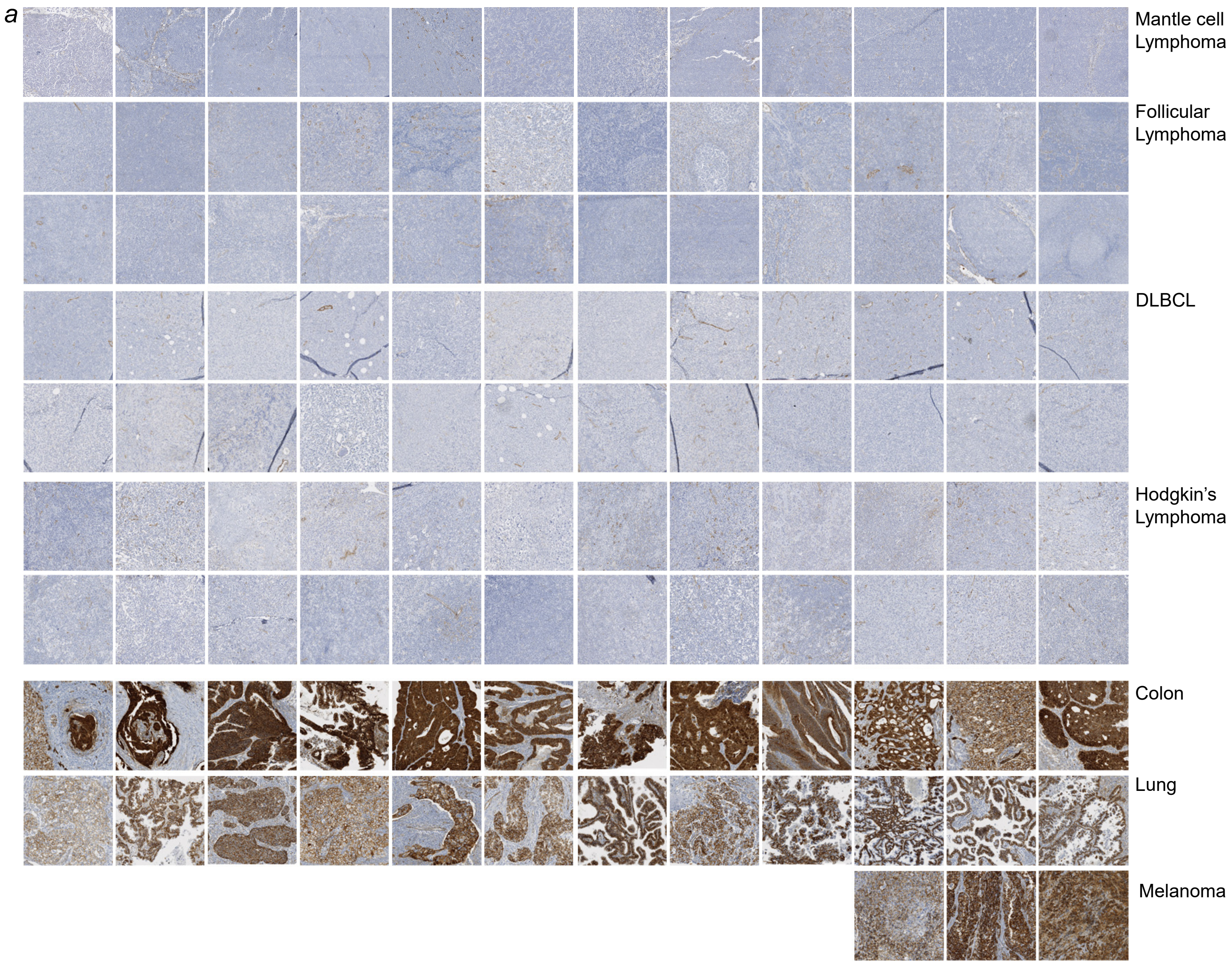


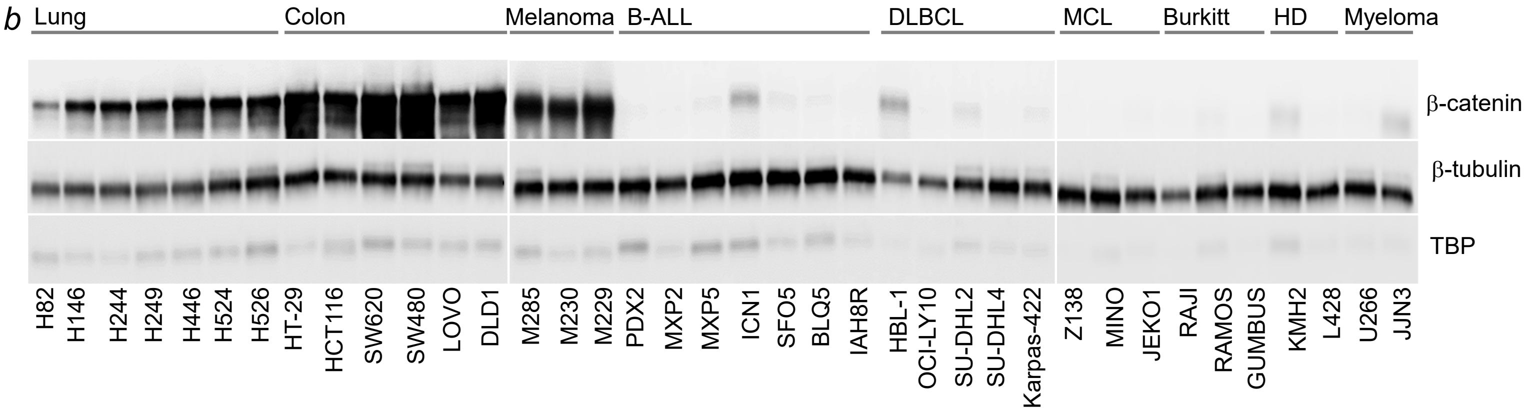


(**a**) β-catenin expression was visualized by immunohistochemistry (brown, counterstain H&E) on tissue microarrays from B-lymphoid malignancies, including mantle cell lymphoma (n=12), follicular lymphoma (n=24), DLBCL (n=24) and Hodgkin's lymphoma (n=24), as well as epithelial cancers, including colon cancer (n=12), lung cancer (n=12) and malignant melanoma (n=3).

(**b**) Western blot analysis of β-catenin expression in cytoplasmic fractions of epithelial cancers, including lung and colon cancer, malignant melanoma, as well as B-lymphoid malignancies, including B-ALL, diffuse large B-cell lymphoma (DLBCL), mantle cell lymphoma (MCL), Burkitt's, Hodgkin's disease (HD) and multiple myeloma cell lines. β-tubulin and TBP were used to indicate purity of cytoplasmic and nuclear fractions, respectively. Western blots of the nuclear fractions from the same cell lysates are shown in **Figure 1g**.

**Figure S3:** *Genetic accumulation of β-catenin suppresses B-cell development in vivo*

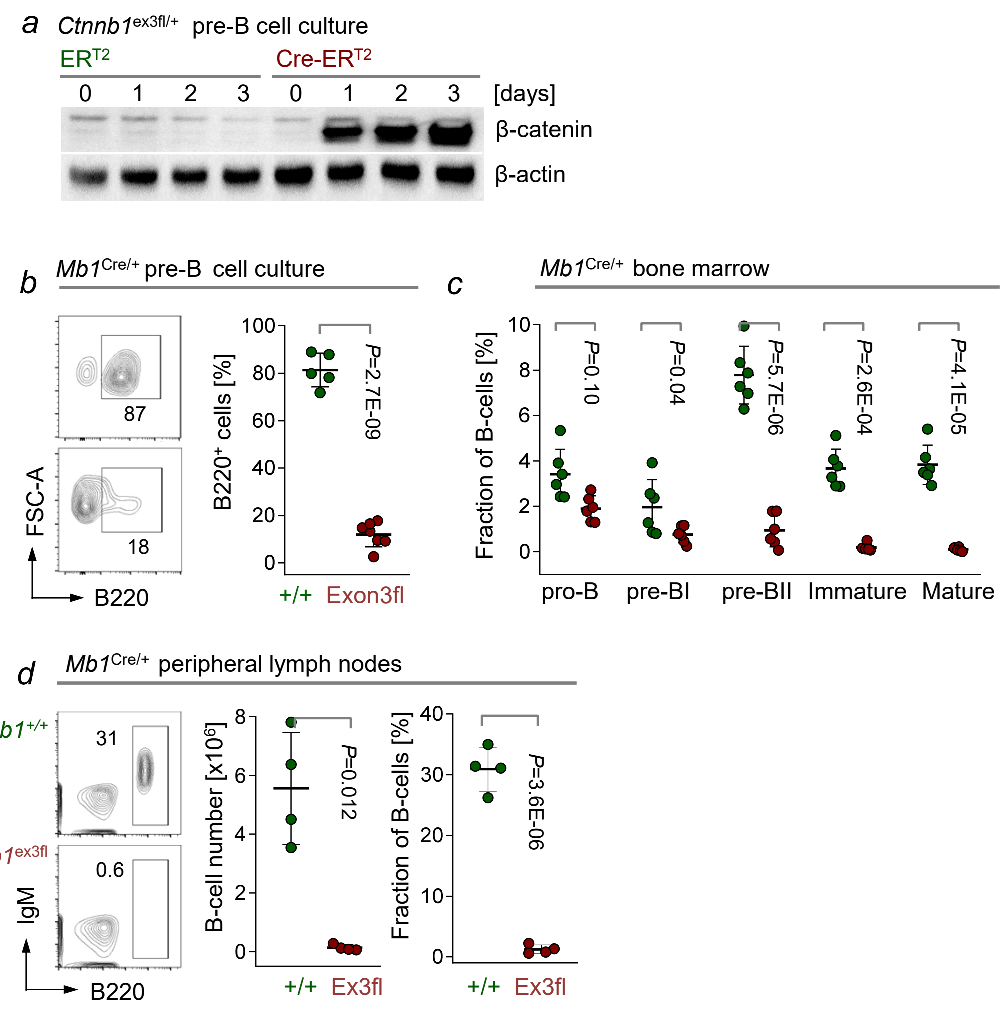


(**a**) Bone marrow pre-B cells from *Ctnnb1*$^{ex3fl/+}$ mice were transduced with 4-hydroxy-tamoxifen (4-OHT)-inducible Cre-ER$^{T2}$ or ER$^{T2}$ constructs. Upon addition of 4-OHT, activation of Cre leads to excision of GSK3β-phosphorylation sites[30], preventing GSK3β-mediated degradation of β-catenin. Western blot analysis was performed to visualize β-catenin accumulation at the times indicated following 4-OHT addition.

(**b**) *Ctnnb1*$^{ex3fl/+}$ mice were crossed with *Mb1*$^{Cre/+}$ for B-cell-specific excision of GSK3β-phosphorylation sites. *In vitro* differentiation of hematopoietic stem cells from the bone marrow of *Mb1*$^{Cre/+}$ *Ctnnb1*$^{ex3fl/+}$ and *Mb1*$^{Cre/+}$ *Ctnnb1*$^{+/+}$ control mice into pro-B and pre-B cells was studied in the presence of IL7. The frequencies of B220$^{+}$ B cells were measured by FACS 7-11 days after removal of Flt3L and SCF. Data shown represent a pool of 5 independent experiments.

(**c-d**) B-cell development in the bone marrow and peripheral lymphoid organs of *Mb1*$^{Cre/+}$ *Ctnnb1*$^{ex3fl/+}$ (red) and *Mb1*$^{Cre/+}$ *Ctnnb1*$^{+/+}$ (green) mice was studied by flow cytometry. (**c**) Relative fractions (%) of B-cell precursor subsets in the bone marrow of the mice are shown for both genotypes. Bone marrow B-cell precursors were distinguished as pro-B cells (CD43$^{+}$ B220$^{low}$ IgM$^{-}$ BP1$^{-}$), pre-BI cells (CD43$^{+}$ B220$^{low}$ IgM$^{-}$ BP1$^{+}$), pre-BII cells (CD43$^{-}$ B220$^{low}$, IgM$^{-}$), immature B cells (CD43$^{-}$ B220$^{low}$ IgM$^{+}$) and mature B cells (CD43$^{-}$ B220$^{high}$ IgM$^{+}$). (**d**) Representative FACS plots, absolute numbers and fractions (%) of B-cells in the peripheral lymph nodes of *Mb1*$^{Cre/+}$ *Ctnnb1*$^{ex3fl/+}$ (red) and *Mb1*$^{Cre/+}$ *Ctnnb1*$^{+/+}$ (green) mice are shown.

**Figure S4:** *B-lymphoid-specific β-catenin-Ikaros factor complexes*

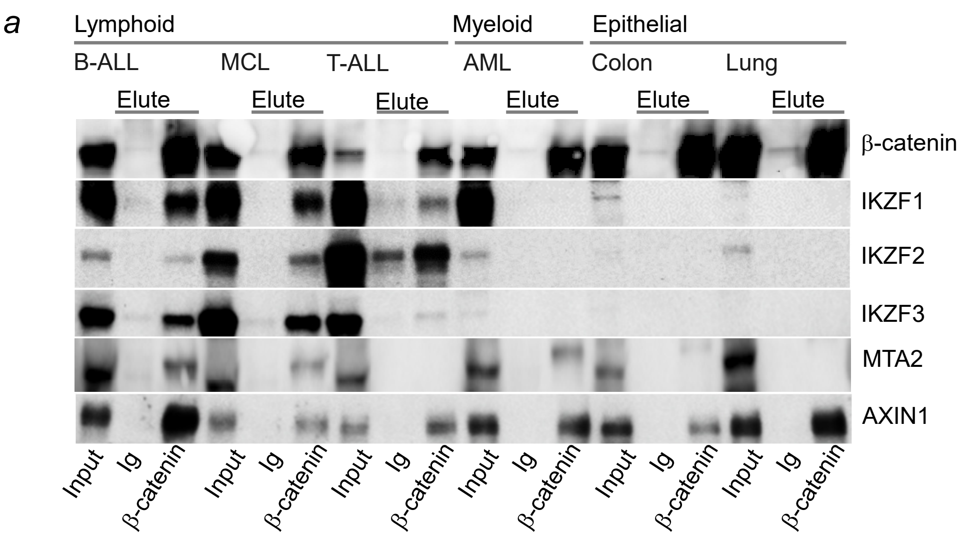


(**a**) B-ALL (MXP2), B-cell lymphoma (JEKO), T-ALL (KOPT-K), AML (MOLM13), colon cancer (SW480) and lung cancer (H446) cell lines were transduced with Tet-3G transactivator and Tre-3G for doxycycline-inducible expression of β-catenin. Co-IP experiments with antibodies against β-catenin or control IgG were performed for input cell lysates, β-catenin-binding of Ikaros factors IKZF1, IKZF2, IKZF3 and NuRD complex component MTA2, as well as AXIN1 (positive control), shown by Western blot.

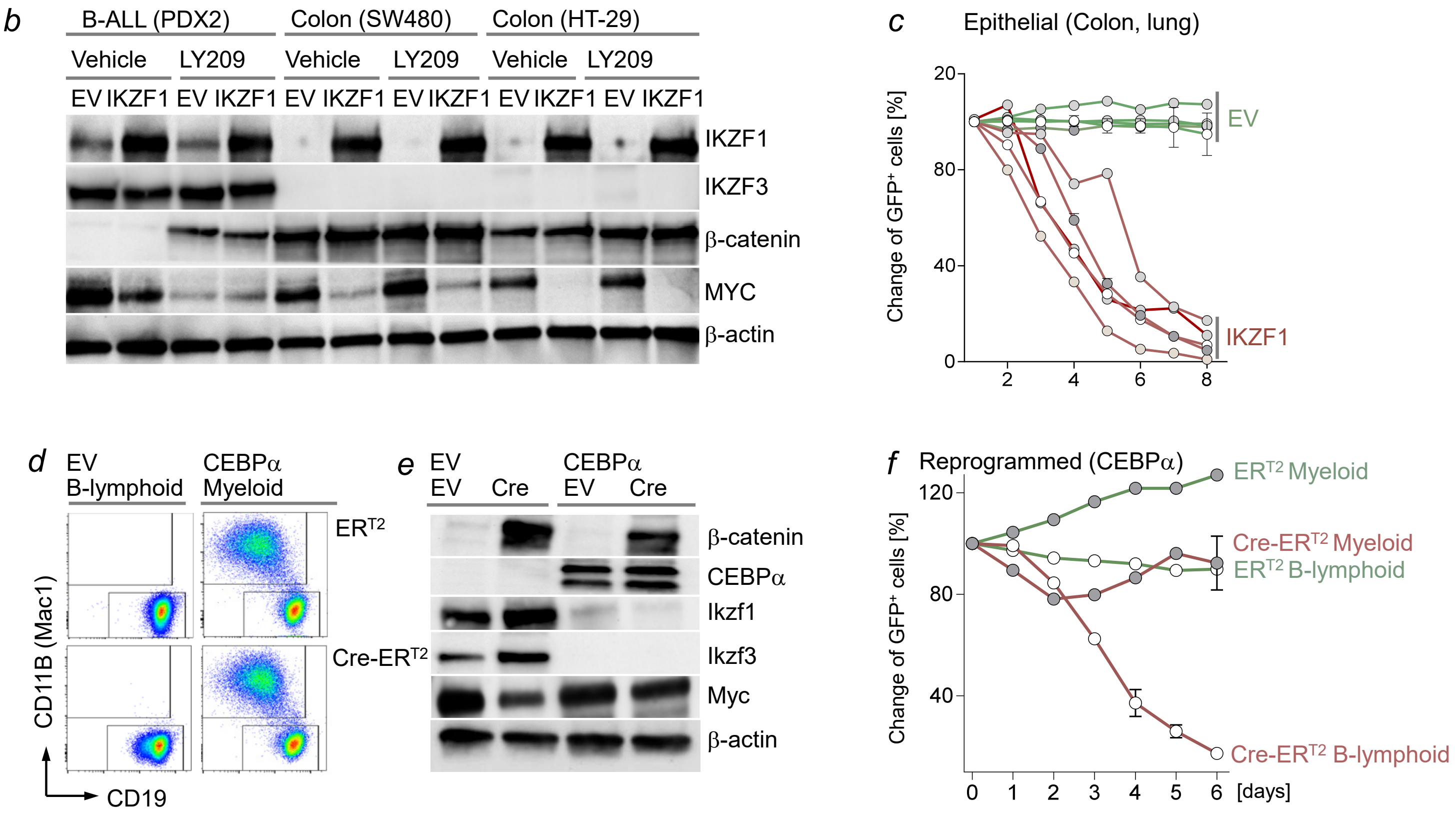


(**b-c**) B-ALL (PDX2), colon (n=3) and lung (n=3) cancer cell lines were transduced with Tet-3G transactivator and Tre-3G for doxycycline-inducible expression of GFP-tagged IKZF1 or EV. (**b**) B-ALL (PDX2) and colon (SW480, HT-29) cancer cell lines were treated with doxycycline for 2 days to induce IKZF1 expression and 16 hours with the GSK3β small molecule inhibitor LY2090314 to accumulate β-catenin. Western blot was performed to study the MYC levels in relation to IKZF1 and β-catenin expression. (**c**) Expression of GFP-tagged IKZF1 or empty vector were induced by addition of doxycycline. $GFP^+$ cells were monitored by flow cytometry. Changes in frequencies of $GFP^+$ cells were normalized to controls. Representative data of 2 independent experiments (triplicates).

(**d-f**) *Ctnnb1*$^{ex3fl/+}$ B-ALL (*BCR-ABL1*) cells were transduced with Tet-3G transactivator and Tre3G for doxycycline-inducible expression of the myeloid transcription factor CEBPα or empty vector (EV). B-ALL cells carrying inducible CEBPα were subsequently transduced with GFP-tagged Cre-$ER^{T2}$ or $ER^{T2}$ vectors for excision of GSK3β phosphorylation sites. CEBPα-driven myeloid reprogramming was induced upon addition of doxycycline. (**d**) Flow cytometry analysis was performed to identify myeloid ($Mac1^+$) and B-lymphoid ($CD19^+$) cells two days after doxycycline treatment. (**e**) Western blot analysis to measure CEBPα, Ikzf1, Ikzf3 and Myc levels following β-catenin accumulation in B-ALL after CEBPα myeloid reprogramming (CEBPα) or EV conditions. (**f**) Changes in frequencies of $GFP^+$ cells were monitored by FACS for 6 days after 4-OHT mediated activation of Cre and accumulation of β-catenin. Data shown is a representative of three independent experiments with three replicates each.

**Figure S5:** *Interactions between Ikaros factors and β-catenin in transcriptional regulation*

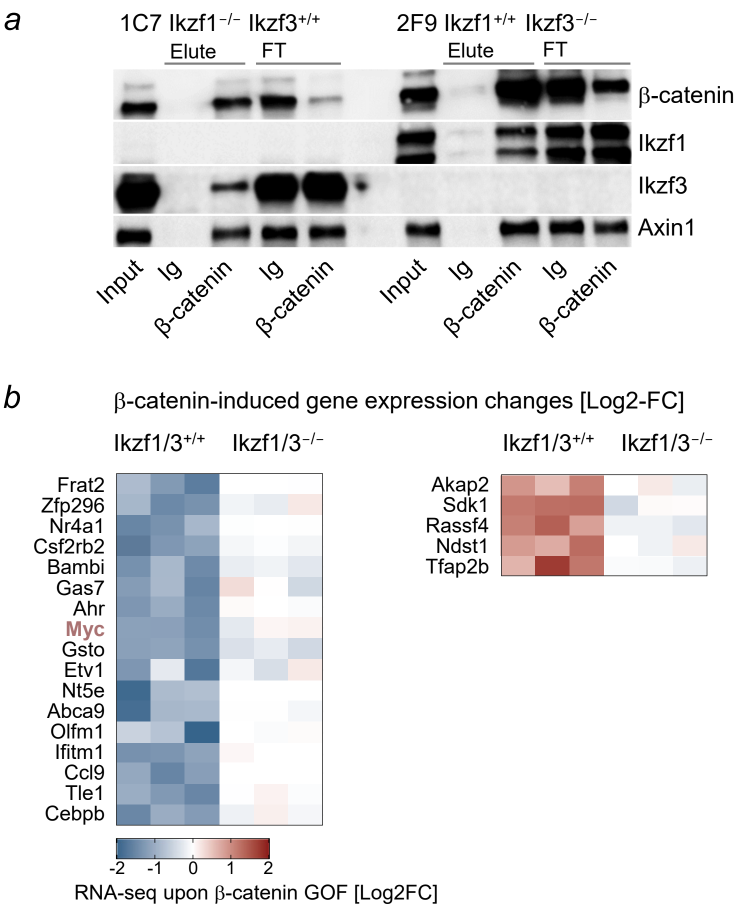


(**a**) To assess whether Ikaros factors can only bind to β-catenin as Ikzf1/Ikzf3 heterodimers (i.e. both Ikzf1 and Ikzf3 are required for binding), we engineered CRISPR-mediated deletion of *Ikzf1* (clone 1C7 Ikzf1$^{-/-}$ Ikzf3$^{+/+}$) and *Ikzf3* (clone 2F9 Ikzf1$^{+/+}$ Ikzf3$^{-/-}$) in *Ctnnb1*$^{ex3fl/+}$ B-ALL cells carrying 4-OHT-inducible Cre-ER$^{T2}$. Co-IP for β-catenin was performed in cells with single deletion of *Ikzf1* or *Ikzf3*. Western blot analysis for β-catenin, Ikzf1, Ikzf3 and Axin1 in whole cell lysates (input), proteins bound (elute) and flow through (FT) to antibodies against β-catenin or control (Ig). Binding of Ikzf1 alone and Ikzf3 alone to β-catenin remained intact.

(**b**) Gene expression changes induced by β-catenin accumulation in the presence and absence of deletion of both Ikaros factors (Ikzf1/3$^{-/-}$) are shown as heatmap. Genes that are repressed (left heatmap) or activated (right heatmap) by β-catenin and how gene expression changes were affected by Ikaros-deletion are shown.

(**c**) Effects of Ikaros factor-deletion (Ikzf1/3$^{-/-}$) on expression of β-catenin target genes (y-axis, log2 fold change) vs. β-catenin binding (x-axis, log2 fold change) are shown. (**d**) Correlation of gene expression changes (y-axis, log2 fold change) with changes in active enhancer regions, H3K27ac signal (x-axis, log2 fold change) upon loss of *Ikzf1* and *Ikzf3*. Absence of Ikaros-factors allowed *de-novo* binding of β-catenin (gained, red) and activation of extra-lineage genes such as Tead1, Tbx19, Lmo1, Lhx2 while lymphoid-specific genes such as Blk1, Bach2, Foxo1 lost binding of β-catenin (lost, blue), H3K27ac and were silenced. Note prominent increased β-catenin binding and increased H3K27ac marks at Myc-BENC superenhancer regions (**d**) upon Ikaros-deletion, consistent with increased Myc expression (**c**).

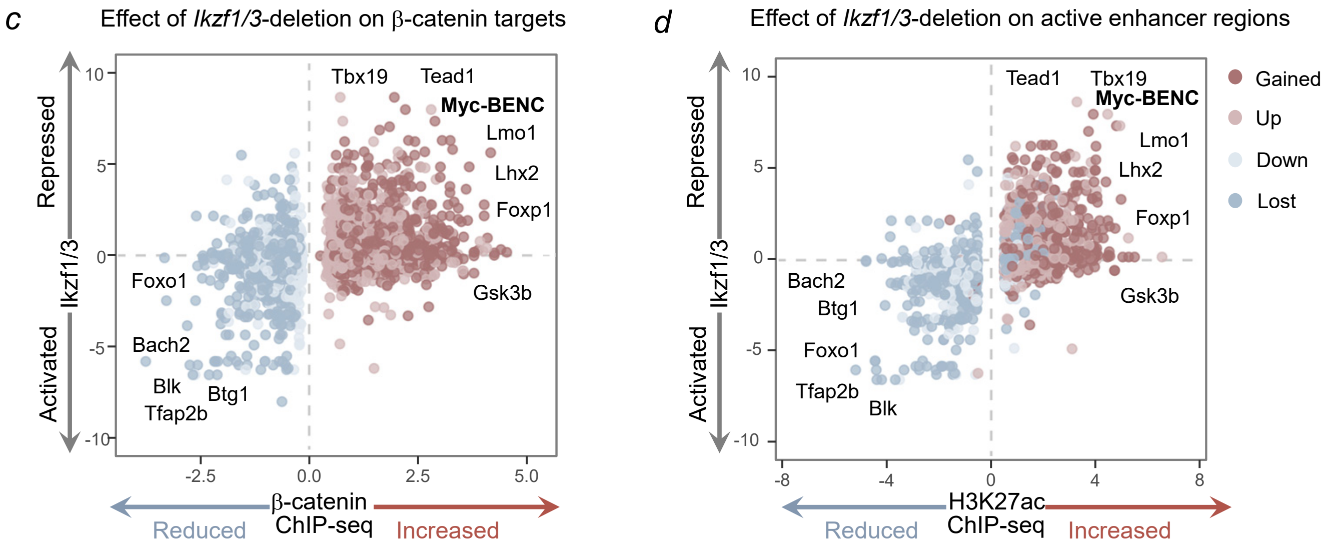

**Figure S6:** *Lenalidomide-induced degradation of Ikaros factors relieves β-catenin-mediated repression of MYC*

*a*

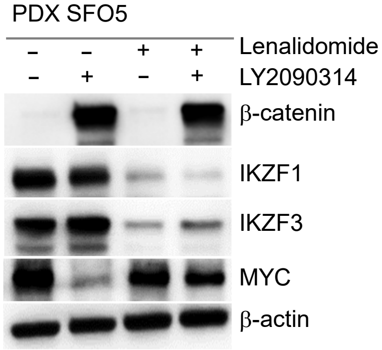


(**a**) Patient derived B-ALL xenografts (PDX, SFO5) were treated with lenalidomide (0.5 μM) to induce CRBN-CRL4-mediated degradation of IKZF1 and IKZF3 Ikaros factors. SFO5 cells were treated with the GSK3β-inhibitor LY2090314 (20 nM) to accumulate β-catenin. Western blot analysis was performed for β-catenin, IKZF1, IKZF3, MYC and β-actin.

*b*

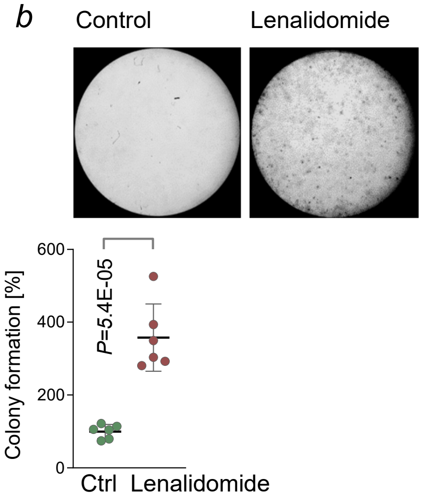


(**b**) Human B-ALL xenograft cells (SFO5) were treated with lenalidomide (0.5 μM) or vehicle for 2 days and plated for colony formation experiments. Representative images and normalized counts (setting mean of vehicle controls as 100%) from two independent experiments (triplicates) are shown.

**Figure S7:** *Ikaros factors profoundly impact β-catenin-binding and β-catenin-mediated gene expression but not vice versa*

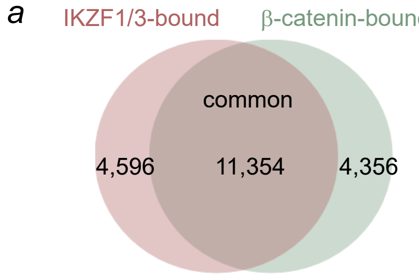


ChIP-seq analysis was performed for Ikzf1, Ikzf3 and β-catenin in *Ctnnb1*$^{ex3fl/+}$ B-ALL cells upon β-catenin accumulation and deletion of Ikaros factors. (**a**) Venn diagram shows the number of regions only bound by β-catenin (4,356), Ikaros factors only (4,596) or both (11,354). Of 15,710 β-catenin peaks, 11,354 (72.2%) were also bound by Ikaros factors.

(**b**) Changes in global distribution of Ikzf1 and Ikzf3 peaks upon β-catenin accumulation. 87% of Ikzf1 and Ikzf3 peaks remained unchanged upon β-catenin accumulation. Color coding for light green box (β-catenin baseline), red box (β-catenin accumulation) and dark green box (Ikzf1/3 baseline).

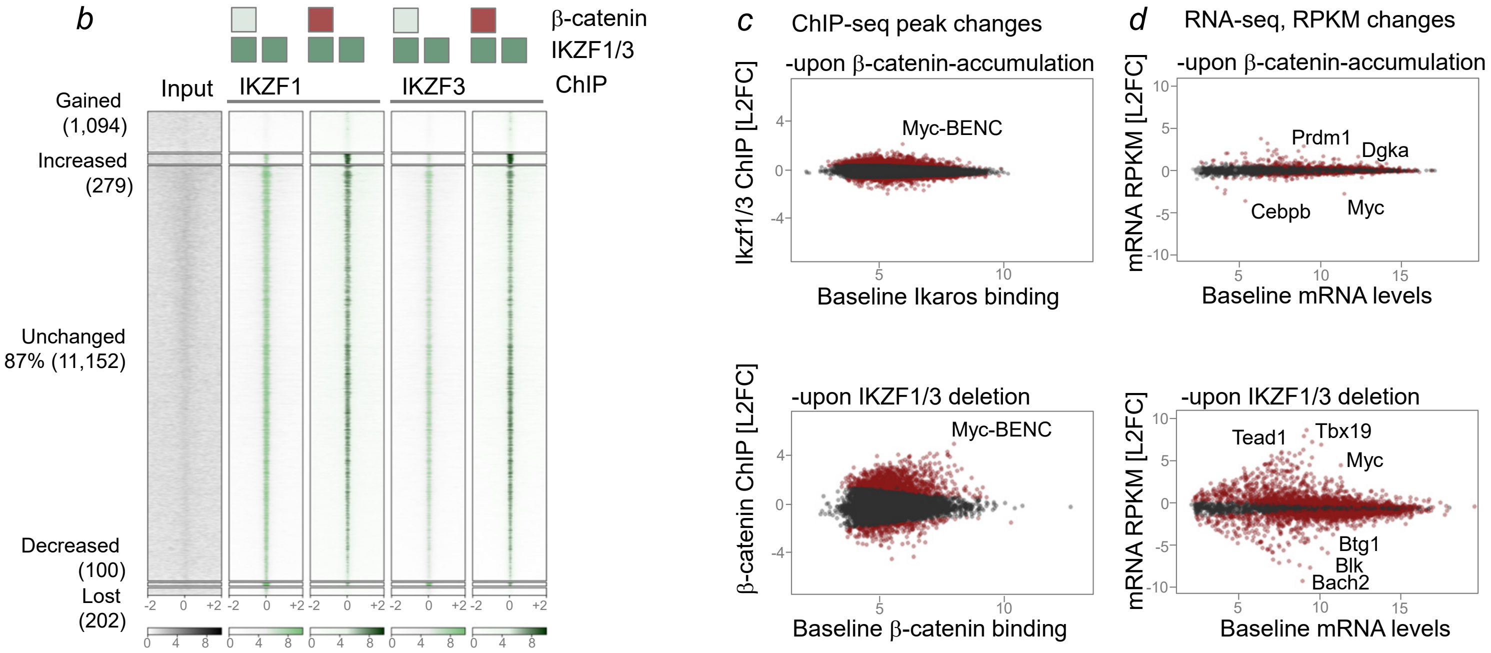


(**c**) Effects of β-catenin accumulation on Ikaros-factor binding (*top*) and effects of Ikaros factor deletion on β-catenin binding (*bottom*) are shown as dot plots for individual ChIP-seq peaks. For each peak, x-axes denote baseline ChIP-seq signals and y-axes show log2-fold changes for Ikaros binding upon β-catenin accumulation (*top*) and β-catenin-binding upon Ikaros deletion (*bottom*). (**d**) Likewise, effects of β-catenin accumulation (*top*) or Ikaros factor deletion (*bottom*) on mRNA levels are shown. Baseline levels for each gene are shown on the x-axes, log2-fold changes denoted on the y-axis for each target gene.

**Figure S8:** *Ikaros factors compete with TCF7 family transcription factors for binding to β-catenin*

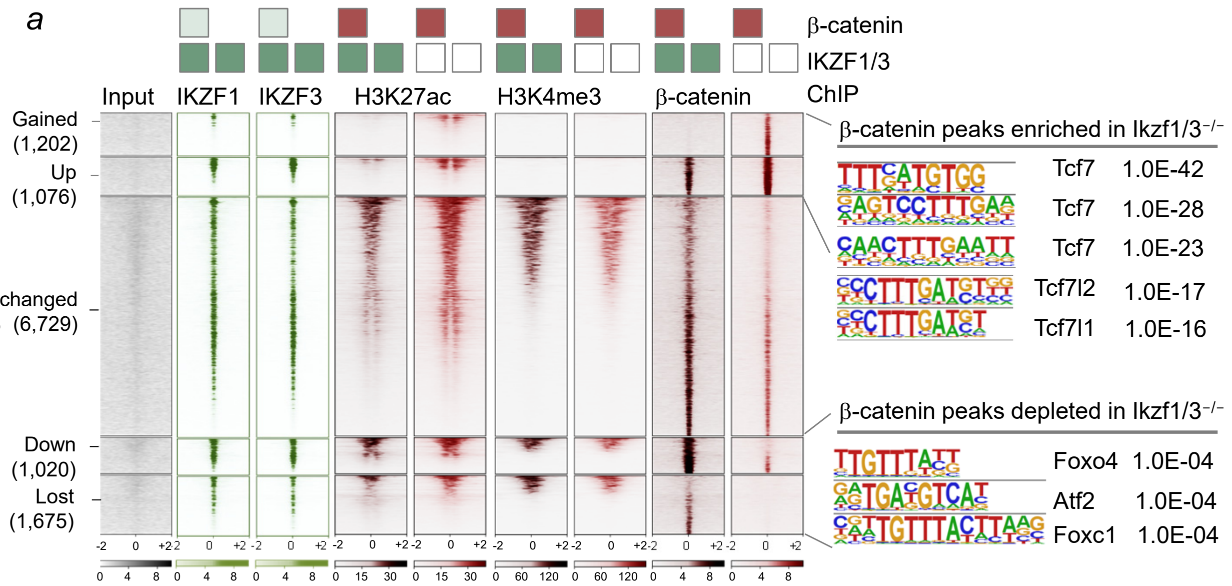


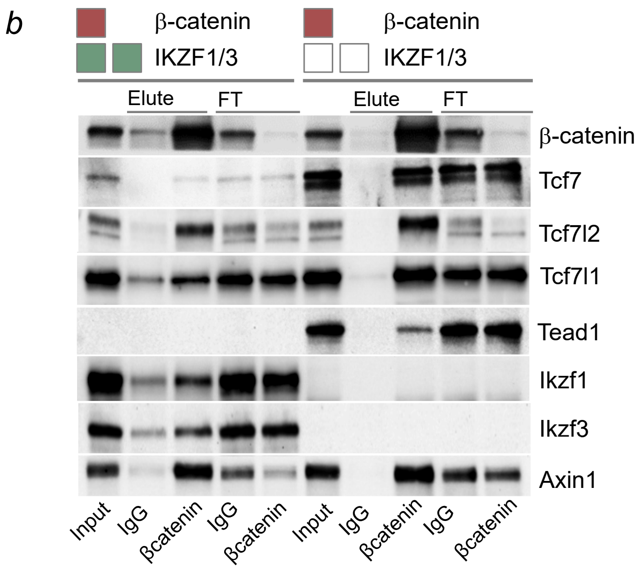


(**a**) ChIP-seq analysis to study genome-wide distribution of β-catenin peaks, colocalization with Ikzf1 and Ikzf3 Ikaros factors as well as H3K4me3 and H3K27ac histone marks. Changes of Ikaros and β-catenin peaks as well as H3K4me3 and H3K27ac histone marks were assessed in the presence and absence of Ikaros deletion (empty boxes) and inducible accumulation of β-catenin (red boxes).

*Ikzf1* and *Ikzf3* deletion enabled binding of β-catenin to inactive enhancers and their subsequent activation (gained H3K27ac). Regions that gained β-catenin binding (n=1,202) are bound by Ikzf1 and Ikzf3 Ikaros factors (*P*=2.5E-62). Deletion of Ikaros factors resulted in redistribution of b-catenin to canonical Tcf7 (*P*=1.0E-42), Tcf7l2 (*P*=1.0E-17) and Tcf7l1 (*P*=1.0E-16) motifs.

(**b**) β-catenin interacting proteins were studied in the presence or absence of Ikzf1 and Ikzf3 Ikaros factors by Co-IP. Western blot analysis was performed in whole cell lysates (input), proteins bound (elute) and flow through (FT) after Co-IP with antibodies against β-catenin or control Ig antibodies. Co-IP was performed under conditions of β-catenin accumulation (red box) and in the presence Ikaros factor deletion (empty boxes) or Ikaros baseline levels (green boxes). Deletion of Ikaros factors enabled binding of β-catenin to Tcf7 and increased interactions with Tcf7l2 and Tcf7l1.

(**c**) Scenario of transcription factor complexes with β-catenin in epithelial cells and B-lymphoid cells: β-catenin pairs with TCF7/TCF7L1/TCF7L2 factors for transcriptional activation of Myc at Wnt responsive elements (WRE) and epithelial enhancer regions (*top*). In B-cells, Ikaros factors outcompete TCF7 to bind to β-catenin. Thereby, Ikaros factors and β-catenin cooperate for effective recruitment of repressive NuRD complexes to lymphoid BENC enhancer regions of Myc, resulting in transcriptional repression of Myc (*bottom left*). Loss of Ikaros factors (Ikzf1 and Ikzf3) enables interactions between β-catenin and TCF7-family factors to restore transcriptional activation of Myc (*bottom right*).

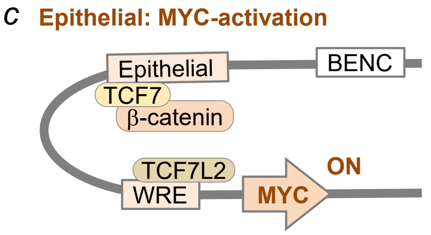


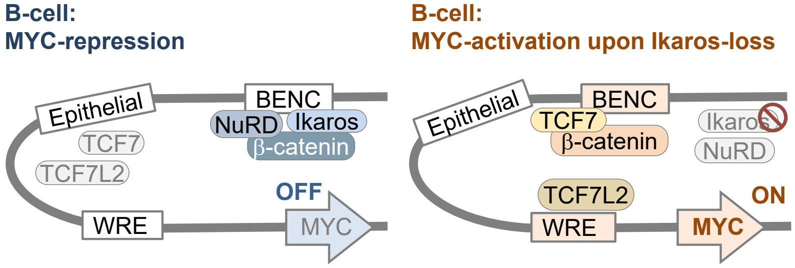

**Figure S9:** *β-catenin enables Ikaros-mediated tumor suppression*

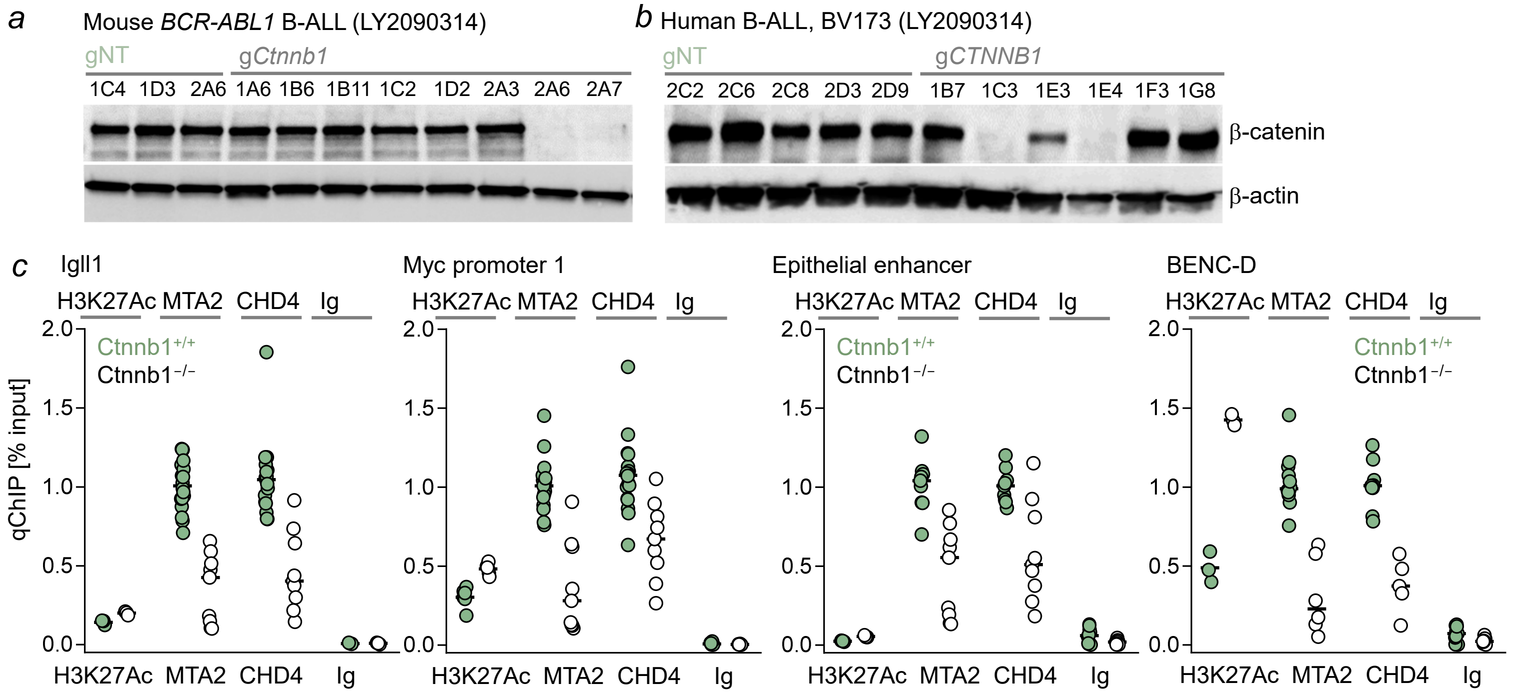


(**a**) Mouse and (**b**) human (BV173) B-ALL cells were gene-edited with guides targeting *Ctnnb1* (g*Ctnnb1*) or non-targeting controls (gNTC). Multiple single-cell clones were selected based on evidence for successful deletion of β-catenin in cells treated with GSK3β-inhibitor LY2090314 to force β-catenin accumulation (Western blot). (**c**) ChIP-qPCR was performed for H3K27ac histone marks, reflecting enhancer activity, and recruitment of NuRD complex components (MTA2 and CHD4) at the *Igll1* promoter (Ikaros target gene), the *Myc* promoter, epithelial Myc enhancer regions as well as the lymphoid BENC Myc enhancer region. For each region, H3K27ac, MTA2 and CHD4 ChIP was performed for B-ALL cells carrying β-catenin deletion (empty circles) or intact β-catenin (light green circles). Data shown represents a pool of four independent experiments.

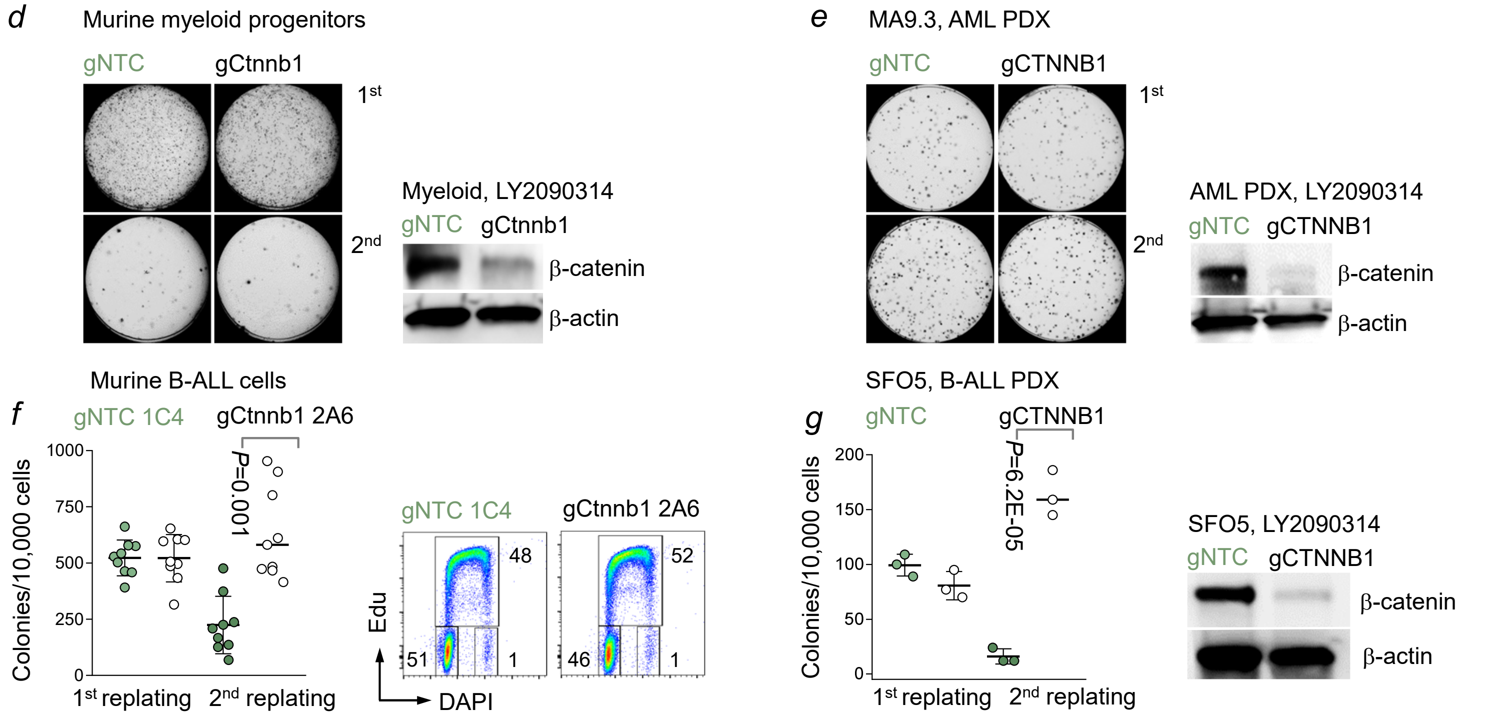


(**d**) Murine myeloid progenitor cells with deletion of β-catenin (gCtnnb1) or non-targeting control (gNTC) were plated in primary (1st) and secondary replatings (2nd) for colony forming assays. Representative images from primary and secondary colonies are shown. Western blot was performed to validate β-catenin loss (representative of two independent experiments). (**e**) Human AML xenografts were edited with guides targeting CTNNB1 (g*CTNNB1*) or non-targeting control (gNTC) and serially plated on methylcellulose medium. Representative images from primary (1st) and secondary (2nd) colonies from two independent experiments are shown. β-catenin deletion was validated by Western blot. (**f**) Mouse B-ALL cells with deletion of β-catenin (clone 2A6) or non-targeting control (clone 1C4) were plated on methylcellulose and 7 days later secondary plating was performed. Number of primary (1st) and secondary (2nd) colonies from three independent experiments are shown (Images, **Figure 5i**). Cell cycle phases were studied by Edu incorporation and DAPI staining (n=2). (**g**) Human B-ALL xenografts (SFO5) with *CTNNB1* deletion (gCTNNB1) or non-targeting control (gNTC) were compared in a serial plating assay. Number of colonies in primary (1st) and secondary (2nd) plating and Western blot for validation of β-catenin-deletion are shown.

**Figure S10:** *β-catenin and Ikaros factors target the BENC-C superenhancer region of MYC*

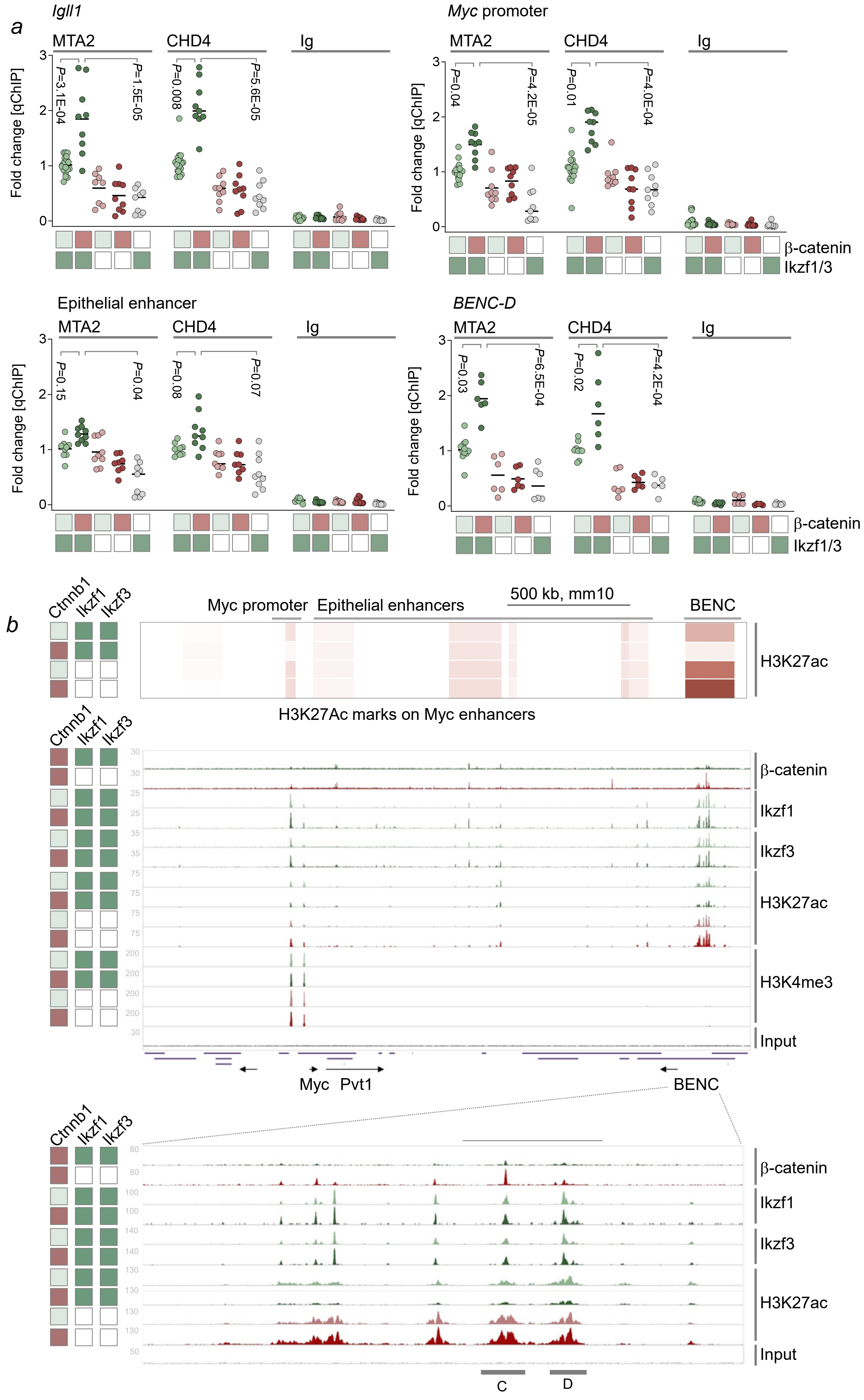

**Legend to Figure S10:** *β-catenin and Ikaros factors target the BENC-C enhancer region of MYC and are both are required for NuRD complex recruitment.*

(**a**) ChIP-qPCR was performed for NuRD complex components (MTA2 and CHD4) at the *Igll1* promoter (positive control as known Ikaros and NuRD complex target gene), the *Myc* promoter, epithelial Myc enhancer regions as well as lymphoid BENC *Myc* enhancer regions. For each region, MTA2 and CHD4 ChIP was performed for B-ALL cells with induced β-catenin accumulation (red boxes), β-catenin deletion (empty boxes) or intact β-catenin (light green boxes), as well as deletion of Ikaros factors (empty boxes) or intact Ikaros factors (dark green boxes). Data shown represent a pool of 6 independent experiments.

(**b**) ChIP-seq analysis for β-catenin, Ikaros factors Ikzf1 and Ikzf3, histone marks H3K27ac and H3K4me3 is shown for the Myc locus, including upstream *Myc* promoter regions and long-range transcriptional enhancers of *Myc* in B-ALL cells from *Ctnnb1*$^{ex3fl/+}$ mice.

Heat map of H3K27ac distribution marking active enhancer regions, shows that most of the H3K27ac enhancer activity is concentrated in lymphoid blood enhancer cluster (BENC) regions in B-ALL cells (*top*). Peak density plots show colocalization of β-catenin, Ikzf1 and Ikzf3 peaks and their concentration at the BENC enhancer regions (*middle*). Close-up view of ChIP-seq peaks of β-catenin, Ikzf1, Ikzf3, H3K27ac at BENC enhancer elements elements C and D in B-ALL cells with accumulation of β-catenin (red boxes), β-catenin baseline (light green boxes), Ikaros factor deletion (empty boxes) or Ikaros baseline (dark green boxes) is shown (*bottom*).

Ikaros factors and β-catenin show marked enrichment at BENC-C and BENC-D regions. While accumulation of β-catenin depleted H3K27ac marks at BENC-C and -D enhancer regions, β-catenin had the opposite effect and increased BENC-C enhancer activity and H3K27ac signals when Ikaros factors (*Ikzf1* and *Ikzf3*) were deleted ($P$=0.002, **Figure 6B**).

**Figure S11:** *Repurposing clinically approved GSK3β-inhibitors for refractory B-cell malignancies*

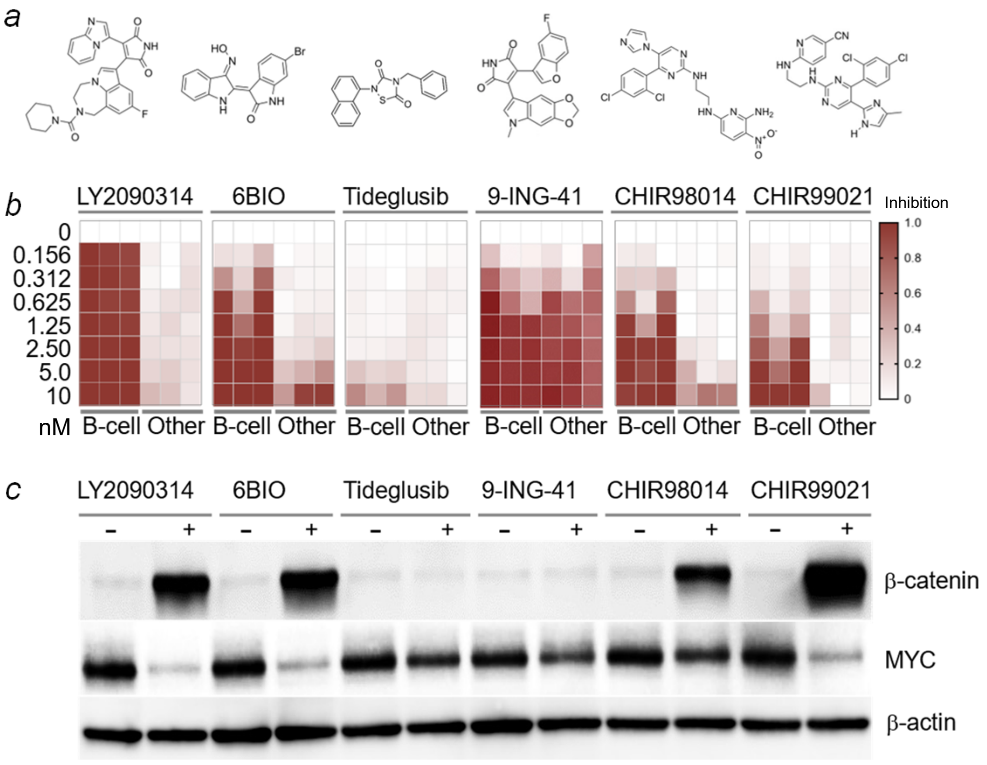


Responses to GSK3β small molecule inhibitors were assessed in three B-cell leukemia (B-cell) cell lines and each one myeloid, colon and lung cancer cell line (Other).

(**a**) Chemical structures of tested compounds are shown. (**b**) Drug responses are shown as a heatmap for LY2090314, 6-bromo-indirubin 3'-oxime (6BIO), Tideglusib, 9-ING-41, CHIR98014 and CHIR99021 in B-cell lines (PDX2, BV173, LAX2) vs. other cell lines (THP1, SW620, H82) at the indicated concentrations.

(**c**) B-ALL (PDX2) cells were treated with indicated GSKβ inhibitors for 16 hours. Changes in protein levels of β-catenin and Myc in relation to β-actin levels were shown by Western blot.

*d*

| Compound | Indication | NCT Identifier | Outcome | Adverse effects |
|---|---|---|---|---|
| LY2090314 | Gastrointestinal cancer, pancreatic carcinoma | NCT01287520, NCT01214603, NCT01632306 | Phase 1 and 2, no clinical responses, favorable safety profile, MTD 80 mg i.v. | Lymphopenia, anemia, diarrhea |
| 9-ING-41 (Elraglusib) | Advanced sarcomas, salivary gland carcinoma, pancreatic carcinoma, melanoma | NCT03678883, NCT05239182, NCT04239092, NCT05077800, NCT04906876, NCT03678883, NCT05116800, NCT04218071, NCT04832438, NCT05010629 | Phase 1 and 2, no clinical responses, IND withdrawn, favorable safety profile | |
| Tideglusib | Alzheimer's disease, myotonic dystrophy, progressive supranuclear palsy, tooth repair (dentin) | NCT01350362, NCT00948259 | Phase 1 and 2, no clinical responses, favorable safety profile | |
| AZD1080 | Alzheimer's disease, Parkinson | Trial in Sweden[1], PMID: 23410232 | Phase 1, favorable PK/PD and safety profile | |
| CHIR99021 (Laduviglusib) | NK-cell infusion for ovarian cancer, solid tumors, hearing loss | NCT03081780, NCT03213964, NCT03319459, NCT03616223 | Phase 1 and 2, no clinical responses, favorable safety profile | Local infections (ear) |

(**d**) Summary of Phase I and Phase II clinical trials with GSK3β inhibitors for a variety of clinical indications. In a total of 22 clinical trials, all tested small molecule inhibitors achieved favorable safety and PK/PD profiles at micromolar plasma concentrations (Cmax). None of inhibitors achieved clinical responses. Moderate adverse effects included diarrhea, anemia and lymphopenia.
